# Co-adapting but not static predators facilitate prey adaptation to a fluctuating environment

**DOI:** 10.64898/2025.12.19.695435

**Authors:** Lou Guyot, Aryan Ramachandran, Luis-Miguel Chevin

## Abstract

Understanding how between-species interactions influence adaptation and population persistence in the face of environmental change is one of the greatest challenges for modern ecology and evolution, with important implications for conservation, and other applied fields. As predation is ubiquitous and may have strong demographic and selective impacts, it is likely to alter how prey respond to a changing environment. However, whether and how adaptation in prey depends on the way selection operates on predators remains little understood. We investigate this question by modeling the evolution of a prey’s trait whose optimum phenotype for fitness changes with the abiotic environment, and which also influences predation via its match with a trait of a predator species. We first show that when coevolutionary processes explicitly emerge from interactions among individuals, maladaptation in prey is not proportional to the difference between their optimum phenotype and that of predators, as assumed in most coevolutionary models. In a fluctuating environment, how well the prey track their moving optimum crucially depends on whether and how the predator evolves. Adaptive tracking of the optimum in prey is facilitated if predators track the same optimum, but hampered if predators have a fixed optimum, or cannot evolve. With eco-evolutionary dynamics, phenotypic mismatch reduces the population size of predators, thus decreasing the strength of predatory selection, but the qualitative influence of predation on prey maladaptation remains otherwise similar. Our findings highlight the importance of the evolutionary context of predators for their impacts on prey, and challenge conservation strategies for prey based on removing their predators.

**Significance statement:** Species in the wild face the dual challenge of adapting to their changing physical environment and coping with detrimental interactions with other species, including predators. While evolution by natural selection may jointly overcome both these challenges, the outcome of this process depends on whether the selective forces from predation and environmental change reinforce or oppose each other. Here, we show that the influence of coevolving predators on prey adaptation to a changing environment strongly depends on the selective scenario for the predators. Adaptation in prey is facilitated if predators also adapt to the changing environment, but hampered if selection favors a constant phenotype in predators. Our results challenge the relevance of predator removal as a conservation strategy to preserve prey.

## Introduction

When attempting to understand and predict eco-evolutionary responses to changing environments, such as ongoing anthropogenic global changes, one of the main challenges lies in accounting for the combined influences of abiotic and biotic environmental factors (1–3). A prominent question in that regard is: To what extent is adaptation of a focal species to a changing abiotic environment influenced by its ecological interactions with other species that do, or do not, adapt to the same changing environment (1–3)? If ecological interactions turn out to be critical to environmental responses in each species, then the research program of species distribution models (SDM, e.g. (4)), including mechanistic niche modelling (5), would be compromised (requiring the use of multi-species joint SDM (6)), as would be studies of thermal tolerance/performance curves (7–9), and the characterization of evolutionary potential solely with respect to the abiotic environment (10–12). On more fundamental grounds, we need to understand whether (and to what extent) the current variation in performance and fitness among abiotic environments in a given species, as measured by environmental tolerance or performance curves (7–9), results only from past exposures to these environments in the evolutionary history of that species, or is also influenced by its history of ecological interactions with other species across these abiotic environments (13).

Predation, and more broadly speaking consumer-resource or victim-exploiter interactions, have long been recognized as a crucial determinant of ecological (14) and evolutionary (15) dynamics, with cascading effects on foodweb structure and resource acquisition strategies. From the standpoint of a prey species that needs to adapt to a changing environment, a critical question is: How, and how much, can this adaptation be altered by the presence of a predator species? From a conservation perspective, removing predators may intuitively seem like a simple way to increase the abundance of a threatened prey population. However in practice, such manipulations have revealed that the effects of predation on prey are more complex than a mere reduction in their abundance (16, 17). This is especially true when a changing environment imposes further selective pressures on prey. Recent experimental (18) and theoretical (19–21) results have demonstrated that predation may in some cases help prey adapt to a changing environment, thereby counteracting its negative effects on population abundance, and sometimes even facilitating prey persistence (22). In particular, two theoretical studies have found that predation can increase the persistence time of prey under directional (linear) environmental change, notably by allowing them to more closely track environmentally driven changes in their optimum phenotype for fitness (20, 22). Similar results have been obtained about harvesting (e.g. fishing, hunting), a form of predation by humans (23, 24). However, these models: (i) did not explore contrasting evolutionary scenarios for predator adaptation to environmental change; (ii) treated the predator-prey interaction through a mean-field approach (assuming that each prey interacts with the mean predator, and vice versa), potentially overlooking important effects that emerge from individual-level processes; and (iii) focused primarily on directional change that eventually favors infinite trait values, rather than on bounded environmental fluctuations that can be sustained indefinitely while keeping phenotypes within a limited range. Natural environments oscillate over different timescales, from daily to yearly (seasonal) to multi-annual (e.g. El Niño–Southern Oscillation, North Atlantic oscillation, …). These cycles can influence natural selection and evolution for organisms with shorter generation times than the environmental period, as demonstrated for Darwin’s Finches with respect to El Niño (25), and for fruit flies with respect to seasonality (26, 27). In addition, virtually any environment exhibits random noise characterized by its variance and autocorrelation structure (28), and the patterns of these fluctuations are currently modified by global change (29).

Here, we study how predators influence prey adaptation and persistence in a temporally fluctuating environment, by combining the theory of adaptive tracking of a moving optimum phenotype (30–33) with a trait-matching model of predator-prey coevolution (34–39). Moving optimum models, where a changing environment causes displacements of the optimum phenotype for fitness, are classically used to investigate adaptation to a changing abiotic environment (reviewed in (33)), and are supported by empirical analyses of long-term datasets from natural populations (40). Trait-matching models of coevolution assume that the strength of the interactions between species (predation, parasitism, competition, mutualism…) is maximized (i.e. stronger competition, higher probability of predation, or of parasitic infection) when the phenotypic values of the interactors match. Trait-matching interaction is a classic assumption in the theoretical literature on coevolution (34–39), and can encompass a range of ecological situations. For predation, the match may involve the same trait in both partners (e.g. preferred habitat or spatial location (41, 42), phenological timing of life history events (43), or body size (42, 44)), or different traits (e.g. visual (45) or chemical (46) detection and recognition, toxin-resistance systems (47, 48)), provided that these traits can be measured in the same unit, or that a conversion can be made between them (e.g. expressing the mismatch in each species as the difference between its phenotype and the one that would maximize the interaction with the other species). While we here rely on this classic trait-matching assumption of coevolutionary theory, we go one step further than most previous coevolutionary models (34–38), by starting from individual-level processes and scaling up to the population level, so as to express the determinants of coevolutionary dynamics from basic demographic parameters of the interaction. We use this model to ask: i) How does divergent selection between the predator and the prey affect their maladaptation in a constant environment? ii) How does predation affect prey adaptation to a changing environment, depending on whether the predator (co)adapts or not? And (iii) To what extent does the answer to these questions depend on how fitness influences population growth in both species, and on the possibility for eco-evolutionary dynamics? Our analysis reveals that individual-level ecological interactions can have unexpected effects at the population level, and that the impact of predation on prey adaptation to a changing environment critically depends on the ecological and evolutionary context for the predators.

## Results

We consider a model (specified in detail in the Methods) where selection on a prey’s trait *x* (with mean phenotype 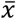) results from a compromise between selective predation on one hand, and adaptation to the non-predatory environment on the other hand, as illustrated in Figure 1. In line with previous coevolutionary theory (34–39), we assume that predation intensity is highest when the prey and predator are phenotypically matched, and denote as *y* the trait of the predator (with mean 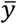). This is modelled though the interaction function 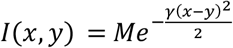 (eq. 9 in the Methods) that relates predation probability to the phenotypes of the prey and its predator, where *M* is the maximum predation probability (when prey and predators are matched), and *γ* is the matching selectivity of the interaction, which determines how much the predation rate is reduced by a given magnitude of mismatch. This predation probability in turn influences the survival of prey and the fertility of predators (blue curves in Fig. 1; eqs. A1 and A4 in the Supplementary Appendix). The associated components of directional selection on the mean phenotypes of prey and predators, termed predatory selection gradients, have directions and magnitudes given by the product of the mean phenotypic mismatch 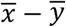 by species-specific strengths of predatory selection, which we denote *γ*_*x*_ and *γ*_*y*_ (eqs. A2, A3, A5 and A6 in the Supplementary Appendix), in line with previous theory (34, 35). For adaptation to the non-predatory environment, we assume that stabilizing selection (with strength *S*_*x*_ in prey and *S*_*y*_ in predators) acts towards an environmentally determined optimum phenotype *θ*_*x*_ for the prey, and *θ*_*y*_ for the predator (yellow curves in Fig. 1). Provided that predatory selection is not too strong relative to stabilizing selection (34–37), the overall fitness function (green curves in Fig. 1) has a displaced optimum with altered peak width (broader in prey, narrower in predators), relative to the case without selective predators.

**Figure 1:**
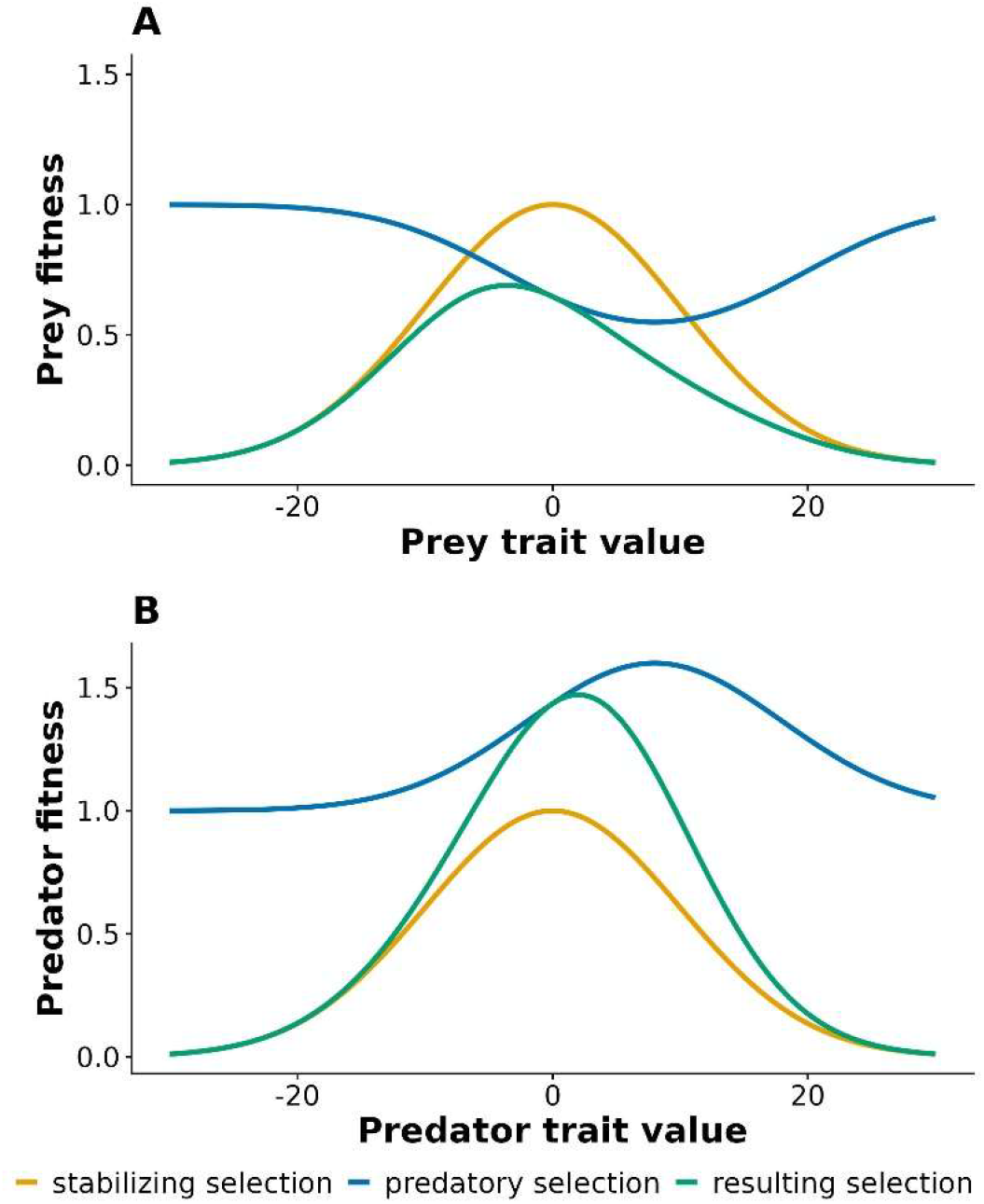
Predatory and stabilizing selection. The components of the fitness function are represented for the prey (A) and the predator (B), for arbitrary values of their optimum phenotype and of the mean phenotype of their interactor, chosen for illustrative purposes. In both cases, stabilizing selection is represented in yellow (using eq. 8), predatory selection in blue (using eq. A1 and eq. A4 from the Supplementary Appendix for A and B, respectively), and the resulting fitness function in green (product of eq. 8 with eq. A1 and eq. A4 for A and B, respectively). Stabilizing selection has strength *S*_*x*_ *= S*_*y*_ *= 0*.*01* and optimum phenotype *θ*_*x*_ *= θ*_*y*_ *= 0*, respectively for prey and predators. For predatory selection, the parameters are the maximum predation probability *M = 0*.*6* and selectivity of predation *γ = 0*.*01*, the population density (*n = 1* in B and *p=1* in A) and mean phenotype of the interacting species (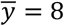 in A and 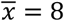 in B), and the baseline reproductive success (*R*_*0*_ *= 1*) and reproductive pay-off of predation for the predator (*α = 1)* in B.

### Coevolution is maximized for intermediate differences between the prey’s and predator’s optimum

As a first step towards understanding how selective predation influences adaptation to the non-predatory environment, we start by analyzing coevolution in a constant environment, with fixed optimum in the prey and the predator (see Table S1 for the corresponding parameter values). When a large number of recombining loci determine the trait values of the predator and prey, we can use results from evolutionary quantitative genetics (34, 35, 49) to predict the equilibrium deviations of the mean phenotypes in predators and prey from their respective optima (eqs. A8 to A18 in Supplementary Appendix), leading to

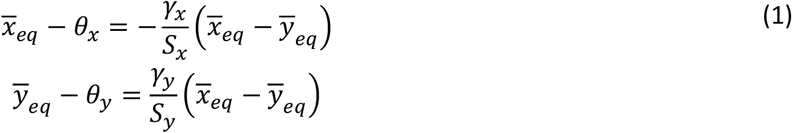

where

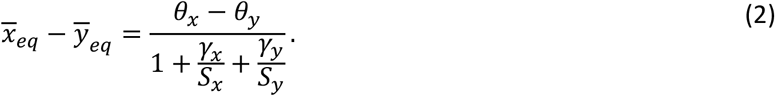

These results – which recapitulate earlier ones from the literature (34, 35) – show that at coevolutionary equilibrium in a constant environment: (i) the prey’s and predator’s mean phenotypes are expected to deviate from their optimum phenotypes (*θ*_*x*_ and *θ*_*y*_ respectively) in proportion to the mean phenotypic divergence between them (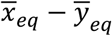 in eq. (1)); (ii) this divergence is in turn proportional to the difference in their optimum phenotypes (*θ*_*x*_ − *θ*_*y*_ in eq. (2)). In particular, if the predator and the prey are selected towards the same optimum (i.e., *θ*_*x*_ *= θ*_*y*_), then trait-matching predatory selection can neither displace their mean phenotypes from each other (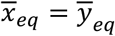 in eq. (2)), nor from this common optimum (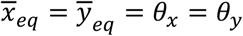, inserting eq. 2 into eq. 1 with *θ*_*x*_ *= θ*_*y*_). And for any non-zero difference in optimum *θ*_*x*_ − *θ*_*y*_, the magnitude of predator-prey phenotypic mismatch, and of deviation of each species from its optimum, depend on the ratios of the strengths of predatory selection *γ*_*x*_ and *γ*_*y*_ over stabilizing selection (*S*_*x*_ and *S*_*y*_), through the composite parameter 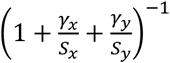.

Previous coevolutionary theory (34–36) treated the strengths of predatory selection *γ*_*x*_ and *γ*_*y*_ as basic population-level parameters, assuming that they were (i) free to vary independently between predators and prey, and (ii) independent of the mean and optimum phenotypes of interactors. However, when building up population responses from individual-level processes, we find that neither of these assumptions holds true, with important consequences for coevolutionary dynamics. Regarding assumption (i), when the predator-prey phenotypic divergence is sufficiently small that 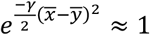 (eq. (A3) and (A6) in Supplementary Appendix), for instance because they have very similar optimum phenotypes, we find that

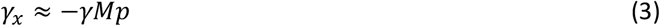

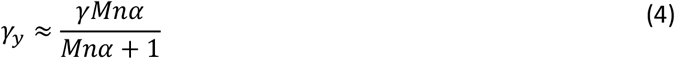

where *α* is the reproductive pay-off of each captured prey for the predator, and *n* and *p* are the population densities of the prey and predator, respectively. Consistent with earlier coevolutionary theory, *γ*_*x*_ and *γ*_*y*_ have opposite sign, because the prey is selected to avoid the predator (*γ*_*x*_ *< 0*), while the predator is selected to chase its prey (*γ*_*y*_ > *0*). But here this sign difference emerges from the individual-level effects of predation on survival and fecundity, rather than being postulated at the population level. Furthermore, eqs. (3-4) show how *γ*_*x*_ and *γ*_*y*_ quantitatively depend on basic parameters of the interaction, as illustrated for *M* and *α* in Fig. S1 (blue lines). In particular, contrary to the assumption from previous coevolutionary theory, *γ*_*x*_ and *γ*_*y*_ are not free to vary independently. Instead, they are coupled by their joint dependency on the product *γM* of the selectivity of the interaction *γ* by the maximum predation probability *M*. The reason for this coupling is that a predation event is necessarily shared among an individual prey and its predator, and so is their interaction function *I*(*x, y*). Interestingly, *γ*_*x*_ and *γ*_*y*_ do not have the same dependency on the selectivity of the interaction: while *γ*_*x*_ increases proportionally to *γ* (eq. 3), *γ*_*y*_ is instead bounded above by *γ*, which it cannot exceed even when both *M* and *α* are large (eq. 4 tends to *γ*_*y*_ ≈ *γ* for large *Mnα*). Lastly, *γ*_*x*_ and *γ*_*y*_ both depend on the population densities of interactors (*n* and *p*), which will prove important for eco-evolutionary dynamics below.

Assumption (ii) on constancy of *γ*_*x*_ and *γ*_*y*_ with respect to the phenotypic mismatch is even more critical. For arbitrary phenotypic divergence between predators and prey, we find that *γ*_*x*_ and *γ*_*y*_ decrease with increasing phenotypic divergence between species (eqs. A3, A6 in the Supplementary Appendix), leading to a feedback between evolution of mean phenotypes and the strengths of predatory selection. However if predatory selection is sufficiently weak relative to stabilizing selection 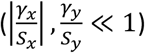, we can approximate the strengths of predatory selection as

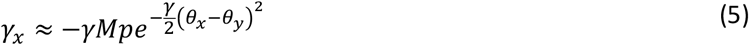

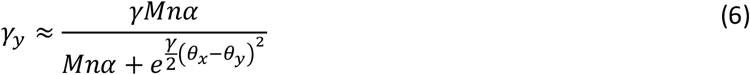

which only depend on basic parameters of the model, not on output dynamical variables such as 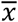 and 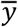. Equations (5-6) show that both *γ*_*x*_ and *γ*_*y*_ tend to 0 as the absolute difference between the optima in predators and prey becomes large (more precisely, for *(θ*_*x*_ − *θ*_*y*_) ≫ *2*/*γ*), as shown in Figure S1 (green lines). This contrasts with the typical assumption in coevolutionary models that *γ*_*x*_ and *γ*_*y*_ are constant with respect to predator-prey divergence and the difference between their optimum phenotypes (34–36). (Note that a weighting similar to eqs. (5, A3) previously appeared in a few studies (38, 39), but was postulated rather than shown to emerge).

Under constant *γ*_*x*_ and *γ*_*y*_, as assumed in previous theory (34–36), deviations of mean phenotypes in prey and predators from their optima increase indefinitely with increasing difference between their optima (eqs. 1-2 combined with eqs. 3-4), but this no longer holds when *γ*_*x*_ and *γ*_*y*_ depend on predator-prey divergence and differences in optimum (as in eqs. 5-6). Under the assumptions leading to eqs. (5-6) (*i*.*e*., 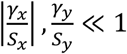, the deviation from the optimum in predators and prey (eqs. 1-2) becomes proportional to 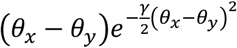, which tends to zero for large |*θ*_*x*_ − *θ*_*y*_|, and reaches a maximum when the absolute difference between the optima in predators and prey is

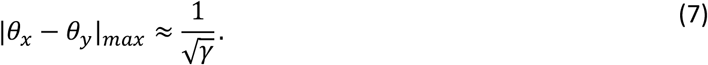

Strikingly, eq. (7) predicts that when eqs. (5-6) approximately hold, the deviation of predators and prey from their optimum are maximized for a specific magnitude of difference in their optimum phenotypes, which only depends on the selectivity *γ* of the interaction, and not on any other parameter of selection. Stronger selectivity causes the maximum to occur at lower differences in optimum.

Individual-based multi-locus simulations on SLiM 4 (50) show that prey and predators evolve away from their optimum (grey lines increasingly deviating from black lines in Fig. 2A), in a way that is well captured by our predictions (colored lines on Fig. 2A). In the following, we quantify the degree of maladaptation for both prey and predators as the deviation of their mean phenotypes from their optimum. The classic linear prediction for prey maladaptation that treats *γ*_*x*_ and *γ*_*y*_ as constant (inserting eqs. 3-4 in eqs. 1-2) works well as long as the difference in optima remains small (blue lines in Fig. 2B). Furthermore, the influence of basic predation parameters on the slopes of these lines is well captured by eqs (3-4) (different values of *M* and *α* across panels of Fig. 2B), illuminating the ecological underpinnings of *γ*_*x*_ and *γ*_*y*_ that had been left implicit in earlier trait-matching theory of coevolution (34–39). When this linear approximation breaks down, the approximation that uses the optimum difference *θ*_*x*_ − *θ*_*y*_ to compute *γ*_*x*_ and *γ*_*y*_ (eqs. 5-6) captures the main tendencies qualitatively (green lines in Fig. 2B), and also quantitatively under weak predatory selection (e.g. *M = 0*.*2* in Fig. 2B). In particular, the prediction that maladaptation in prey is maximized for |*θ*_*x*_ − *θ*_*y*_*| = 10* (from eq. 7 with *γ = 0*.*01*) matches simulation results very well in this case (*M = 0*.*2* in Fig. 2B), and also provides a reasonable proxy under other parameter values (dotted lines in Fig. 2B). The more exact prediction that uses the mean phenotypes from simulations to compute *γ*_*x*_ and *γ*_*y*_ (eqs. A3, A6 in the Supplementary Appendix) almost perfectly matches the maladaptation computed from individual-based simulations (orange lines in Fig. 2B; *γ*_*x*_ and *γ*_*y*_ are shown in Fig. S1). As expected from the assumptions underlying the approximations in eqs. (5-6), the green lines more closely match the simulations results (and the accurate predictions from the orange lines) under conditions where predatory selection is weaker and becomes dominated by stabilizing selection (small *M* and *α* in Fig. 2B). The same conclusions can be drawn for predators (Fig. S2). While the approximations in eqs (5-6) rest on the assumption that stabilizing selection is stronger than predatory selection, in practice they perform reasonably well over a range of values of stabilizing selection strength (*S*_*x*_, *S*_*y*_) and selectivity of the interaction *γ* (Fig. S3).

**Figure 2:**
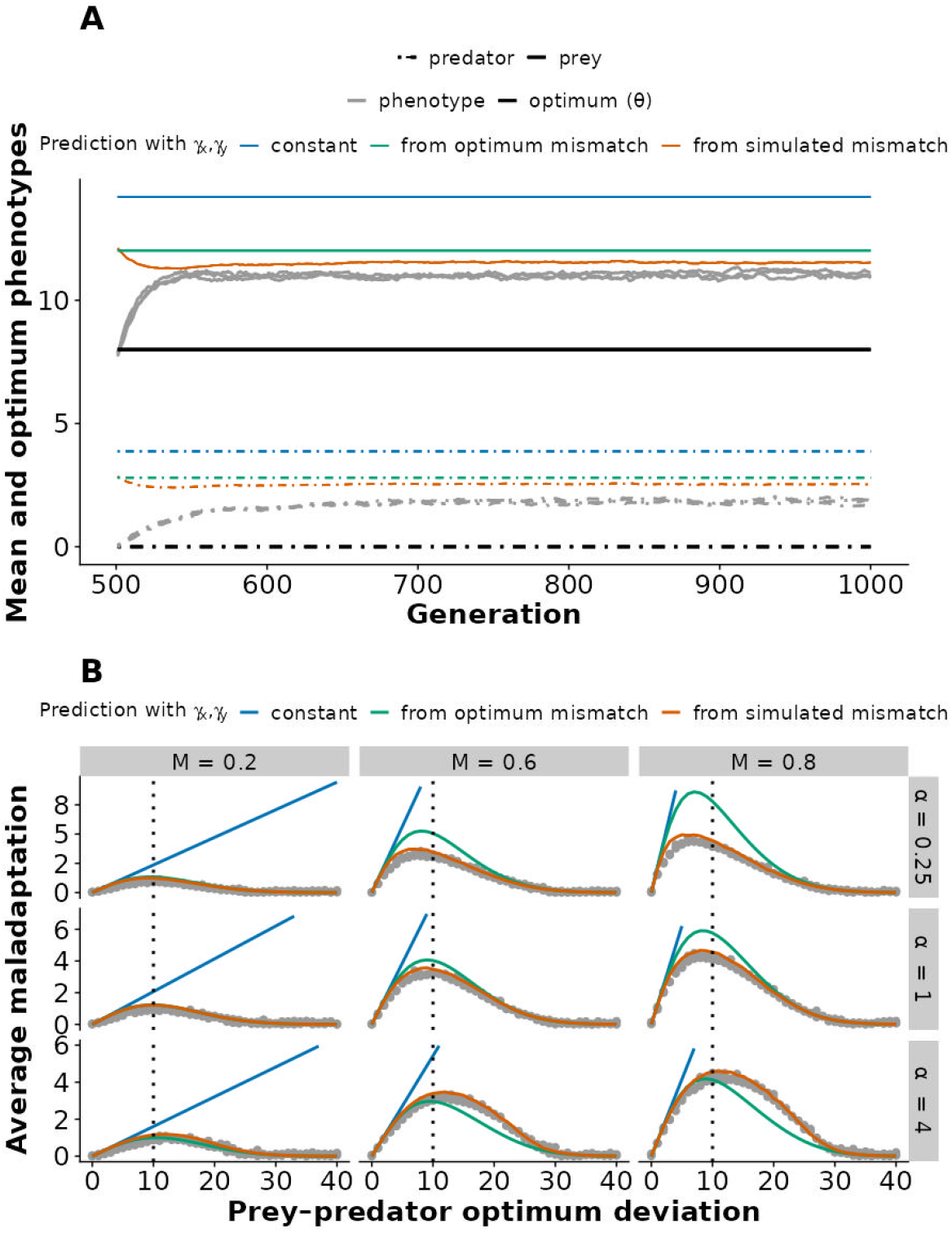
Phenotypic displacement in a constant environment and with constant population sizes. A) The dynamics of mean phenotypes (gray lines) over 500 generations are shown for three replicate simulations of a Wright–Fisher population in a constant environment. Prey are represented with solid lines and predators with dashed lines, and the black lines represent the optimum phenotypes for each species in this environment. The simulations were first run for 500 generations without predation (e.g., assuming the predator initially consumes other prey), allowing each species to reach its optimum phenotype (not shown). Then at generation 500, we introduced phenotype-dependent interaction according to eq. (9), and plotted the changes in the mean phenotypes of predators and prey. Various predictions for the equilibrium mean phenotypes of prey and predators are also represented, using eqs. (1, 2) with *γ*_*x*_ and *γ*_*y*_ given by eqs. (3, 4) (blue), eqs. (5, 6) (green), or eqs. (A3, A6) from the Supplementary Appendix combined with the mean mismatch from three replicate simulations (orange). Note that the orange line does not represent the transient dynamics of the mean phenotype, but only its predicted equilibrium updated by the current values of 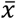 and 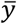 (while the predictions from the blue and green line only depend on basic model parameters). B) The phenotypic deviation of prey from their optimum (gray dots) is plotted against the difference between prey and predator’s optimum phenotypes, for three replicate simulations of Wright–Fisher populations averaged over 500 generations (following a burn-in of 500 generations), across a range of optimum differences (from 0 to 40). Colored lines plot the same approximations as in panel A, and the vertical dotted line indicates 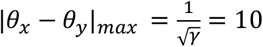 (from eq. 7). Parameter values are *S = 0*.*01, γ = 0*.*01, N* = *P* = 10000; and additionally for panel A: *M* = 0.6, *θ*_*x*_ *= 8*, and *θ*_*y*_ *= 0*, and α = 1.

### Coadapting vs static predators have opposite effects on adaptive tracking in prey

When coevolving populations also experience sustained environmental change, an important question becomes: How well are they able to adapt to these changes by tracking their moving optimum phenotype (30–33), and how is this influenced by coevolution of their interacting species? We here focus on bounded environmental fluctuation where mean phenotypes can reach a stationary distribution in the long run, centering our main argument on deterministic periodic cycles (Fig. 3), but also briefly addressing stationary stochastic fluctuations (Fig. S4).

**Figure 3:**
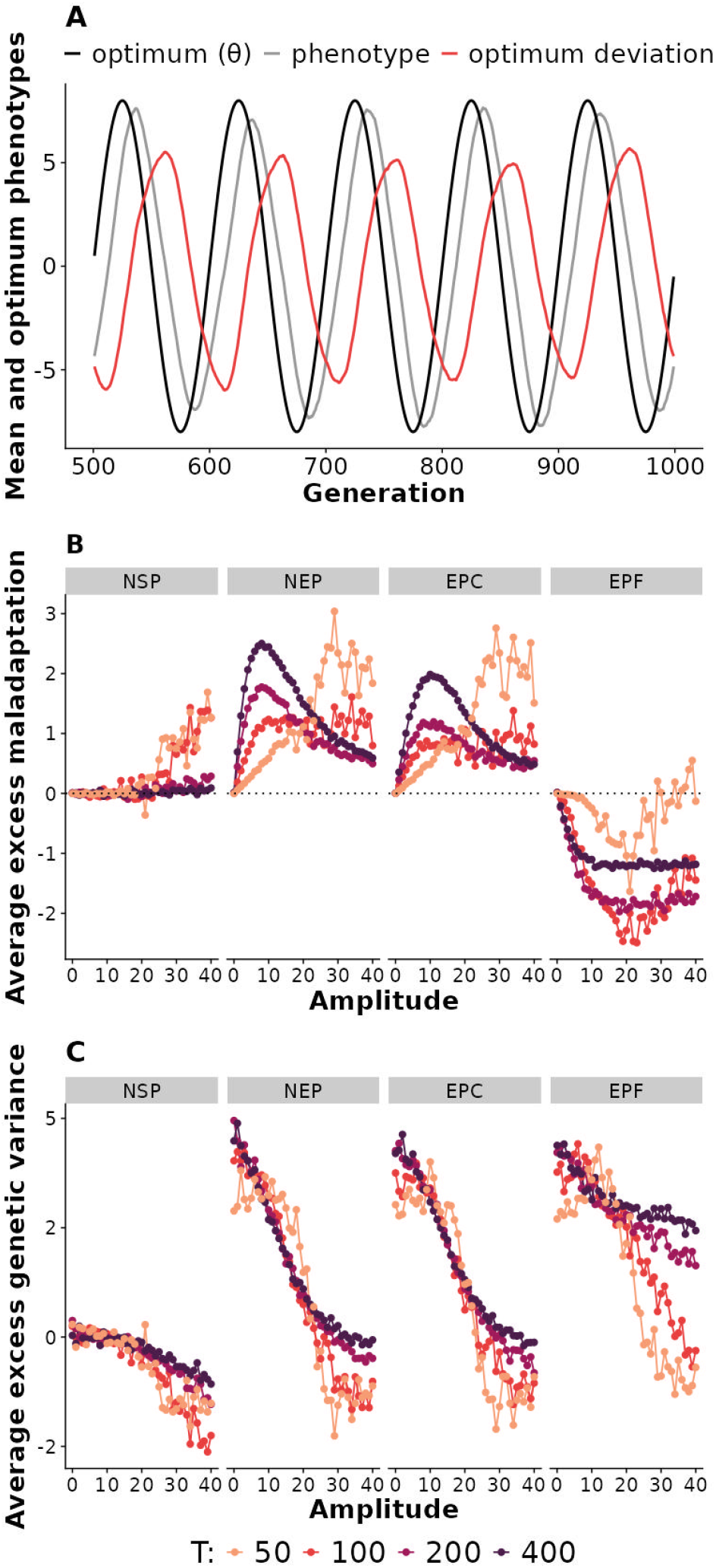
Adaptive tracking in a periodically fluctuating environment with constant population sizes. A) The mean phenotype of prey is shown in gray, for 500 generations of simulations of a Wright–Fisher population adapting to a periodically fluctuating optimum (shown in black), in scenario EPF where a selective predator tracks the same fluctuating optimum as the prey. The deviation of the prey’s mean phenotype from the optimum is also plotted in orange. B) Prey phenotypic deviation from optimum and C) genetic variance, both shown as difference with the no-predator (NP) scenario (denoted as excess maladaptation and excess genetic variance, respectively), are represented against the amplitude of fluctuations in the optimum, for different periods *T* (colors). Four scenarios regarding predators are represented: a non-selective predator (NSP); a non-evolving predator (NEP); an evolving predator with a constant optimum at 0 (EPC); and an evolving predator with the same fluctuating optimum as prey one (EPF). Results are averaged over 100 periods after a burn-in period of 5 periods, under a Wright–Fisher model. Parameter values are *S*_*x*_ *= S*_*y*_ *= 0*.*01, γ = 0*.*01, N = P = 10000, α = 1, M = 0*.*6*; and additionally for panel A: cycle period *T = 100* and amplitude *A = 8*.

When a single species adapts to a periodic environment causing a sine wave in the optimum, both its mean phenotype and deviation from the optimum eventually undergo a sine wave with same period as the optimum, but a lagged phase and reduced amplitude (31, 32, 51). When a prey coevolves with its predator, we also find in our individual-based simulations that the prey’s mean phenotype and deviation from the optimum eventually settle into cycles with the same period as the optimum (Fig. 3A; predators are also shown in Fig. S5), although these cycles may depart from a sine wave to some extent (Fig. S6). We thus summarize the degree of maladaptation in prey as the amplitude 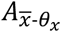 of cycles in their deviations from the optimum. To investigate how (co)evolution of the predator influences maladaptation in the prey, we contrast different scenarios (Table 1): a non-selective predator (NSP); a non-evolving predator (NEP); an evolving predator with a constant optimum (EPC); and finally an evolving predator with a fluctuating optimum that moves like the prey’s optimum (EPF), which we describe as coadaptation. The scenario without any predator (NP) is also used as a baseline for comparison.

**Table 1:** Selective scenario for predators explored in the simulations with a fluctuating environment.

| Scenario | Definition | Specific parameter values |
| --- | --- | --- |
| No-predator (NP) | Prey are in isolation, without predators | $P = p = 0$ |
| Non-selective predator (NSP) | Predators hunt non-selectively, that is, the probability of predation is determined independently from their phenotypic difference with the prey | $I(x, y) = M$ |
| Non-evolving predator (NEP) | Predation is conditioned on phenotypic similarity between predator and prey, but the predator's trait remains constant, at a value corresponding to the mean optimum of the prey | $S_y = B_y = 0$ |
| Evolving predator with constant optimum (EPC) | Predator can evolve, but their optimum phenotype is not affected by the fluctuating environment, instead remaining constant at a value corresponding to the mean optimum of the prey | $B_y = 0$ |
| Evolving predator with fluctuating optimum (EPF) | Predators evolve and their optimum phenotype changes in the same way as that of the prey | $B_y = B_x = 1$ |

In a periodic environment, even a non-selective predator (NSP) can increase maladaptation in prey relative to the no-predator (NP) scenario (left in Fig. 3B and S7A). This effect is mediated by the predator’s impact on genetic variance in prey. Indeed predation imposes a death toll on prey that reduces their effective population size (Fig. 7C), thereby increasing the intensity of random genetic drift, which in turn decreases the genetic variance (left in Fig. 3C, and Fig. 7B). As the genetic variance of traits results from a dynamic equilibrium between mutation, genetic drift, and (fluctuating) selection (52, 53), it also depends on patterns of environmental cycling, being more strongly reduced under large amplitudes and short periods (left in Fig. 3C), which jointly lead to fast changes in the optimum (the average absolute speed over a cycle being 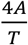). Reduced genetic variance leads to poorer tracking of the optimum under all scenarios (Fig. S7A-B).

A selective predator (scenarios NEP, EPC and EPF) has stronger effects than NSP on adaptive tracking in prey. In particular, a predator that either cannot evolve (NEP), or is not selected to track a fluctuating optimum (EPC), tends to exacerbate prey maladaptation relative to a case with no (NP) or non-selective (NSP) predation (Figs. 3B and S7A, middle). This is especially clear under large periods (*T* = 200 or 400), under which the prey tracks its optimum equally well under NSP and NP (Fig 3B, left), while NEP and EPC lead to large excess maladaptation (Fig. 3B, middle). The reason for this excess maladaptation is that, when the mean phenotype of predators is fixed at the average optimum of the prey (NEP), or forced by stabilizing selection to remain close to this value (EPC), it selectively “pushes” the mean phenotype of the prey farther from its optimum in all environments. The prey then evolve as if it were tracking cycles with a larger amplitude than the optimum (by a factor 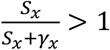, from eq. A20 in the Supplementary Appendix), overshooting its true optimum in all environments. This effect is however non-monotonic with respect to the amplitude of fluctuations, because for large amplitudes, the intensity of predatory selection (*γ*_*x*_ and *γ*_*y*_) decreases owing to the large difference in optimum between the predator and prey (from eq. 5), and so does the influence of coevolution on mean phenotypes (compare NEP and EPC in Fig. 3B to Fig. 2B). The non-linearity that this introduces, together with the changes in genetic variance with the amplitude and period of cycles (Figs. 3C and S7B), complicate the patterns quantitatively, precluding the use of simple analytical approximations to interpret these results. However, the conclusion that a non-coadapting predator (scenarios NEP and EPC) increase maladaptation of a prey in a cycling environment – relative to no or a non-selective predator – holds qualitatively under all conditions we have investigated (Fig. 3B).

The outcome is strikingly different when the selective predator co-adapts to the same cycling optimum as the prey (EPF in Fig. 3B and Fig. S7A). In this case, maladaptation in the prey is reduced relative to case without (NP) or with a non-selective predator (NSP), under all conditions except for very rapid movements in the optimum (large *A* and small *T*, Fig. 3B). The reason for this reduced maladaptation is that a co-adapting predator tracks the optimum with a larger lag than the prey, therefore exerting a net selective push on the prey towards the optimum across the cycle, as under an linear change in the optimum (22). Analytical approximations that assume a constant genetic variance and fixed *γ*_*x*_ (eqs. A19-A31 in Supplementary Appendix) provide further insights into how the different scenarios for predator (co)evolution affect maladaptation of prey in a cycling environment.

In a stationary stochastic (rather than periodic) environment, the differences in coevolutionary outcomes among scenarios of predation (from NSP to EPF) are largely similar to those under periodic cycles - albeit more noisy owing to environmental stochasticity - if we replace the amplitude of cycles by the standard deviation of fluctuations, and their period by a characteristic time of autocorrelation, as specified in the Methods (Fig. S4).

### Eco-evolutionary dynamics reduce the influence of predation on adaptive tracking

When fitness influences population growth in both species, allowing for eco-evolutionary feedbacks, the deviation of the prey from its optimum in a constant environment is smaller than under fixed population sizes (compare Fig. 4A with middle panel in Fig. 2B; also Fig. S8A with Fig. 2B and Fig. S8B-D to Fig. S3A-C). This occurs because the reduced probability of predation caused by the phenotypic mismatch decreases the mean fitness of predators, which now affects their population density *p* (Figs. 4B and S8M-P), in turn leading to a weaker predatory selection *γ*_*x*_ on the prey (from eqs. 3,5; Fig. S8I-L compared to Figs. S1 and S3G-I). Maladaptation in prey therefore becomes non-monotonic with respect to its optimum difference with the predator, even over a range where it would be close to linear with constant population sizes (curved blue line in Fig 4A that account for changes in population size, compared to the straight blue lines in Fig 2B; more parameter values are plotted in Fig S8F-H, to be compared to Fig. S3A-C). Further accounting for the effects of differences in mean and optimum phenotypes on *γ*_*x*_ and *γ*_*y*_ described above (eqs. 3-6) accurately predicts maladaptation in the prey in a constant environment (green and orange lines in Fig. 4A; more parameter values in Fig. S8).

**Figure 4:**
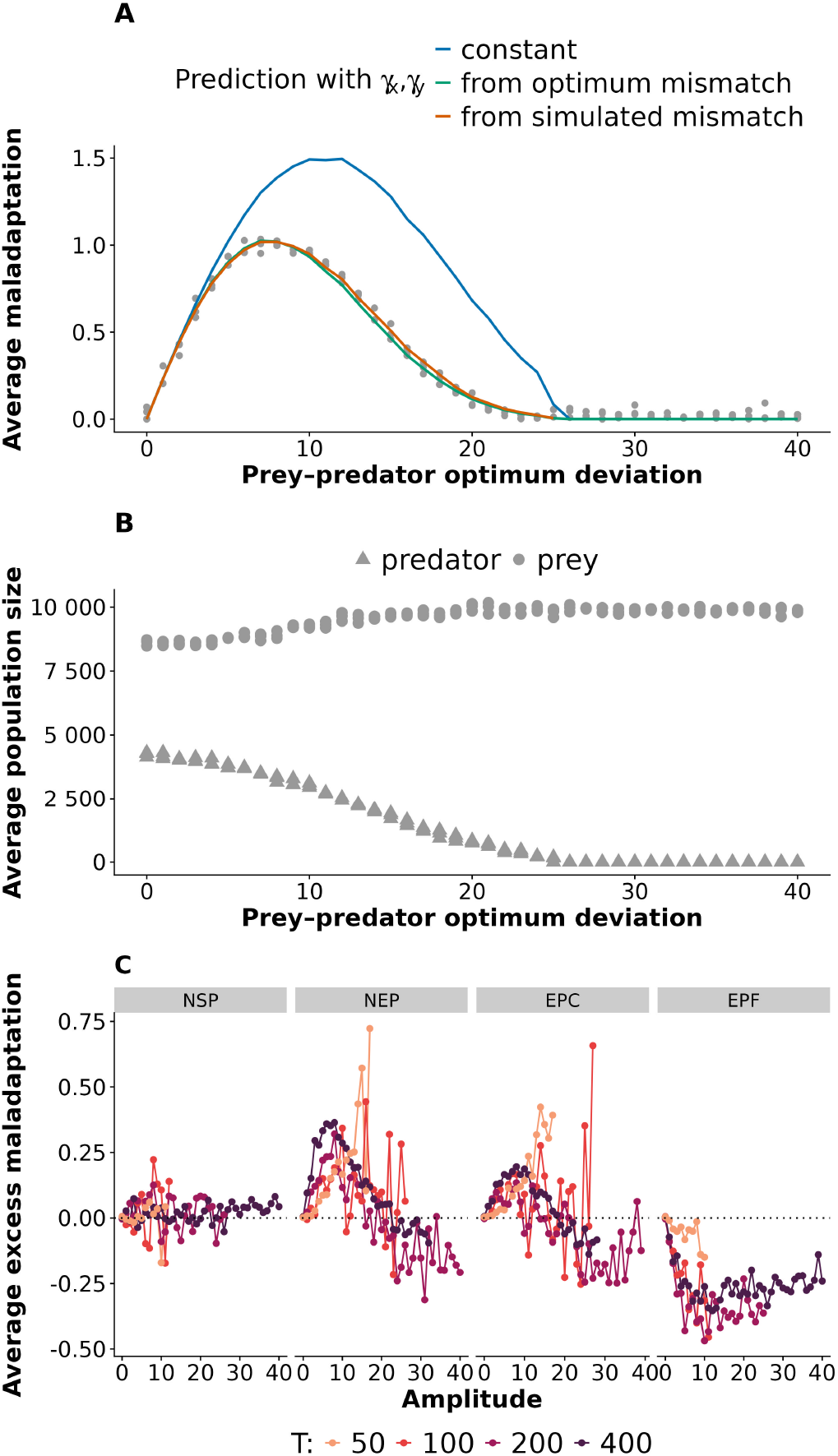
Phenotypic displacement and adaptive tracking with eco-evolutionary feedbacks. The phenotypic deviation from the optimum in prey (A), and the population sizes of prey and predators (B) are plotted against the difference between the prey’s and predator’s optima, for three replicate simulations with a constant environment and eco-evolutionary feedback, averaged over 500 generations (after a burn-in of 500 generations), across a range of optimum differences (from 0 to 40). In panel A, predictions based on different approximations for *γ*_*x*_ and *γ*_*y*_ (colored as in Figure 2) are also shown as colored lines. The effect of a fluctuating environment with eco-evolutionary feedback is shown in C, which plots the average amplitude of phenotypic deviation from the optimum in prey, expressed as a difference with the no-predator scenario (denoted as excess maladaptation), against the amplitude of fluctuations in the optimum. Results are averaged over 100 periods after a burn-in period of 5 periods, in simulations with eco-evolutionary feedback run across a range of amplitudes of fluctuation (from 0 to 40). Missing dot correspond to simulations in which prey or predator populations got extinct. Parameter values are *γ = 0*.*01, S*_x_ *= S*_γ_ *= 0*.*01, M = 0*.*6* and *α = 1*; the period *T* of cycles is specified in the legend (color of lines) in panels B-C.

In a fluctuating environment, the influence of selective predation (scenarios NEP, EPC, EPF) on prey maladaptation is also smaller with than without eco-evolutionary dynamics (Fig. 4C versus Fig 3B), because the reduced (and fluctuating) population density of the predator leads to weaker net selective predation. Nevertheless, the differences between alternative scenarios for selection on predators are qualitatively similar to those without eco-evolutionary feedbacks (Fig. 4C versus Fig. 3B), except that prey maladaptation in NEP and EPC becomes lower than for NP and NSP under large amplitudes of fluctuations. Still, the qualitative conclusions that a co-adapting predator (EPF) always leads to better adaptive tracking in the prey, while a static predator (NEP and EPC) leads to poorer adaptive tracking over a broad parameter range (associated with moderate amplitudes of fluctuations), also holds with evo-evolutionary dynamics.

Extinction of prey and/or predators also becomes important to consider when eco-evolutionary dynamics are allowed. We find that extinction is more likely for the predator than for the prey (Fig. S8M-P), as expected since the former needs the latter to persist, and maladaptation at lower trophic levels cascades up to increase extinction risk at higher trophic levels (54). In a fluctuating environment, rapid environmental change (large *A*/*T* ratio) leads to higher extinction risk (missing dots in Figs. 4C and S9). We find that the parameter range allowing persistence of both the predator and the prey (i.e., periods and amplitudes for which dots are plotted in Fig. S9) is more restricted for NSP and EPF. These scenarios are associated with the most intense predation, either because predation occurs regardless of phenotypes (NSP), or because the predator-prey mismatch remains small (EPF). Beyond their intrinsic importance, extinction events also modify the coevolutionary outcomes relative to the case without eco-evolutionary dynamics, by causing a bias towards conditions compatible with persistence of both the predators and the prey (a demographic constraint on adaptation, *sensu* (55)).

## Discussion

Understanding how ecological interactions alter adaptation to a changing environment is one of the greatest challenges for predicting eco-evolutionary responses to climate change, and other anthropogenic perturbations (1–3). Here we show that the response to this question for a prey species crucially depends on how selection operates on its predator.

### Non-linear effects of predatory selection

Our first key finding is that, when predation is mediated by phenotype matching between the predator and its prey, maladaptation in both species – as measured by their displacement from their optimum phenotype – scales non-linearly with the difference in their optimum phenotypes, decreasing as this difference become large in a constant environment. This differs from the predictions of most previous trait-matching coevolutionary models defined only at the population level (34–37), where the equilibrium maladaptation in each species is proportional to the difference between their optima. The reason for this discrepancy is that, when the contribution of species interactions to fitness is explicitly defined at the level of random encounters and interactions between individuals of these species, the strengths of predatory selection *γ*_*x*_ and *γ*_*y*_ (for prey and predators, respectively) are not constant, but instead decrease with the decreasing probability of interaction, as the mismatch increases. Such a decrease in the strength of interaction-mediated selection was postulated without demonstration in a few studies (38), but was more often ignored altogether (34–36). We here demonstrate that it arises naturally when species interactions have a binary outcome (e.g. predation success or failure) that depends on their phenotypes, over a discrete number of encounters. This decrease in predatory selection *γ*_*x*_ and *γ*_*y*_ with increasing mismatch causes the predatory selection gradients (eqs. A2 and A5 in Supplementary Appendix) to be non-monotonic with respect to predator-prey mismatch, and maximized at an intermediate value, unlike in earlier population-level models (34–37).

This non-monotonicity of selection gradients with respect to the phenotypic mismatch between interactors may lead to coevolutionary tipping points, similar to the evolutionary tipping points described for single species in a directionally changing environment under some fitness functions (56) or life cycles (57). Here, gradually increasing the difference between the optimum phenotypes of the predator and its prey would first increase the predatory selection gradient, but only up to a tipping point (towards a mismatch of 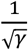, from eq. 7), beyond which further increases would reduce the intensity of predatory selection, causing the traits of both species to evolve back towards their optimum without predation (character release).

### Role of predators in adaptation to changing environments

To investigate how a predator affects adaption to a changing environment in its prey, we focused on adaptive tracking of a stationary fluctuating optimum (either periodic or stochastic), because this imposes sustained selective pressure within a limited phenotypic range (while a purely directional trend in the optimum can only be sustained indefinitely if traits can evolve to infinity). Our results revealed that whether predation helps or hinders adaptation in the prey critically depends on the evolutionary scenario for the predator. A selective predator that either cannot evolve (NEP scenario), or is not selected to track a moving optimum (EPC), may impose substantial phenotypic maladaptation on its prey. In contrast, a predator that needs to track the same optimum as its prey (EPF) can facilitate adaptation in the prey, relative to the case without predation (NP), or to a non-selective predator that does not attack prey based on their phenotypes (NSP).

It has been previously shown that a predator can facilitate adaptation of a prey to a linearly changing environment, if the additional component of directional selection that it causes aligns with that of the changing environment, an effect described as the “selective push” (22). However in a fluctuating environment, the “selective push” for NEP and EPC causes the prey to undergo fluctuations of larger magnitude, and thus to often overshoot their optimum, increasing maladaptation. On the other hand, for EPF the predators lag behind both the optimum and the prey, thereby causing a selective push that brings the prey closer to its optimum in most conditions.

### Changes in genetic variance

While the opposite influence of a co-adapting (EPF) versus non-coadapting (NEP and EPC) predator on prey adaptation is a robust finding that is largely independent of genetic details, other qualitative and quantitative simulation results are partly driven by changes in genetic variance among selective regimes. In particular, the amplitude of cycles in the optimum has a strong influence on genetic variance in the prey, and this effect differs among selective scenarios for predators (Fig. 3B and S7B), limiting the use of analytical results based on the assumption of a constant genetic variance to interpret patterns of maladaptation. Unfortunately, there is a scarceness of analytical predictions for the maintenance of quantitative genetic variance at a balance between mutation, genetic drift, and a periodically fluctuating selection, and the existing theoretical results (without predation) are based on individual-based simulations (52). At least the broad qualitative responses, such as the increase in time-averaged genetic variance with increasing period and amplitude, should be amenable to analytical interpretation when drift can be neglected, but the comparison between scenarios with no versus non-selective predation (NP vs NSP) suggests that genetic drift also plays an important role. While this was not our focus here, future studies should investigate in more detail the maintenance of quantitative genetic variance when a fluctuating environment is combined with (possibly selective) ecological interactions.

### Relevance of model assumptions

We assumed that the predation rate was maximized when a single phenotypic trait matches between the predator and the prey, in line with classic models of coevolution under various types of ecological interactions (34–38). Allowing several traits in the predator to match several traits in the prey would also be possible, and would modify the outcome to some extent, owing to a coevolutionary cost of complexity (58), but would certainly not alter our main conclusions about the influence of different scenarios of adaptation in predators. Alternatively, some studies have considered directional interaction functions, where predation rate increases if the phenotype of the predator exceeds that of the prey (e.g. running speed or arms races (36)). The influence of (possibly co-adapting) predators on prey adaptation to a changing environment is likely to differ in this context, and would be worth investigating in the future.

Our scenarios of adaptation in the predator (NSP, NEP, EPC and EPF) were mostly chosen as limit cases to bound the range of possible outcomes for more realistic ecological settings. However, these scenarios are also ecologically meaningful in some contexts. For instance, EPC (evolving selective predator with a constant optimum) may represent the situation where the higher trophic level mostly needs to track its resource (i.e. the prey), rather than the abiotic environment *per se*. This is likely to be relevant to classic examples of breeding phenology, where e.g. birds are not necessarily strongly constrained by spring temperature, but their laying date needs to match the timing of the peak in caterpillar abundance (their resource), which itself needs to match the timing of budburst in trees (the resource for caterpillars), with the latter being more directly selected by the temperature (to be able to complete the growth and reproductive cycle while avoiding late frosts after bud burst) (59). As this example illustrates, it would be worthwhile extending our model to interaction networks with more species and trophic levels, to understand how the processes we have described cascade through communities and ecosystems, extending previous work that did not include fluctuating environments (38, 60).

### Empirically testing the theory

To our knowledge, only one experimental study, conducted in a constant environment, has directly demonstrated that predators can facilitate prey adaptation to a new environment (18). Beyond this work, few experimental systems have simultaneously considered evolutionary and ecological dynamics in predator–prey interactions. Some experiments using predator–prey systems in constant or fluctuating environments have focused on the traits mediating the interaction (e.g. (61–64)). Others have examined the response of different trophic levels to environmental fluctuations (65), or the impact of invasive predators on prey persistence or extinction (66). However, despite these advances, experimental evidence directly testing how predator coadaptation influences prey evolution under environmental change remains lacking. Evolution experiments using controlled predator–prey systems in changing environments could provide a powerful framework to address this question. Such studies would help determine whether predator evolution can consistently facilitate prey adaptation and persistence, and under which ecological and environmental conditions.

## Conclusions

Our analysis indicates that the influence of a predator on prey adaptation to a fluctuating environment critically depends on how selection acts on the predator, as well as on the magnitude and speed of environmental fluctuations. This has consequences for the interpretation of variation in performance and fitness across environments. Environmental tolerance (or performance) curves are often thought to reveal how organisms evolved in response to their past exposures to relevant abiotic environmental variables (7). However if ecological interactions, such as predation as studied here, or competition (13), strongly modulate adaptive evolution to the abiotic environment, then tolerance curves might as well reflect an evolutionary history of exposure to ecological interactors across environments. This has important consequences not only for our understanding current responses to the abiotic environment, but also for predicting adaptation and population persistence in the face of ongoing and future environmental change, especially considering the rapid pace of current anthropogenic environmental perturbations.

## Methods

### Individual-based model and simulations

We designed an individual-based model of predator-prey interactions and coevolution, which we simulated using SLiM version 4.2.2 (50). The model captures the (co)evolutionary dynamics of a prey species and its predator, exploring different scenarios for: (i) predator presence and evolutionary potential; (ii) environmental fluctuations; and (iii) demography - with either constant population size or eco-evolutionary dynamics. The explored scenarios and parameter values are reported in Tables 1 and S1.

#### Genetics of traits

Adaptation to the (changing) environment and species interactions are assumed to both be mediated by a quantitative trait in each of the species, with *x* the trait of the prey and *y* the trait of the predator. These traits are determined by *L* diploid quantitative trait loci (QTL) with additive effects. Mutations occurs at a rate *µ* per haploid locus, and the effect sizes of these mutations are drawn from a standard normal distribution (hence assuming a mutational variance of 1). The recombination rate between adjacent loci is set to *r*. Genetic parameter values are specified in Table S1.

#### Fitness and selection

Selection on trait *x* in the prey and trait *y* in the predator involves two components: one mediated by adaptation to the (possibly changing) environment (contributing *U*_*x*_ and *U*_*y*_ to fitness), and another mediated by species interactions, i.e. predation (contributing *V*_*x*_ and *V*_*y*_). These components act multiplicatively on fitness, such that the total fitness for trait *z* (where *z = x* for prey and *z = y* for predators) is *W*_*z*_ *= U*_*z*_*V*_*z*_.

The selection component caused by environmental adaptation is modeled by letting traits be under stabilizing selection for an optimum phenotype that may change with the environment. Specifically, we use the classic assumption of a Gaussian fitness peak,

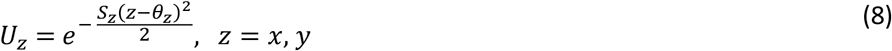

where *θ*_*z*_ is the optimum, and *S*_*z*_ is the strength of stabilizing selection.

The selection component caused by predation emerges from random encounters between predators and prey, followed by phenotype-dependent predation upon each encounter. Each prey is assumed to encounter predators randomly with a probability that depends on their population density. This is modeled by drawing, for each prey, the number of predators it encounters from a Poisson distribution with parameter equal to the predator density 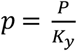 , where *P* is the population size of predators, and *K*_*y*_ their carrying capacity (defined below). Hence *p* is used as a proxy for the proportion of the total area (e.g. number of patches) occupied by the predator. At each encounter, a predator is randomly sampled from the population, and the probability of successful predation is obtained from an interaction function *I*(*x, y*) that depends on the phenotypes of both interactors. Specifically, we follow earlier literature (34–37) in assuming that the predation probability is highest when the phenotypes of predators and prey are matched. This is modelled using a Gaussian function,

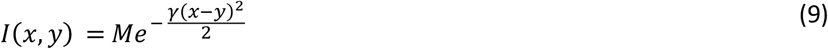

where *M* is the maximum probability of successful predation when traits of the predator and prey match, and *γ* tunes the matching selectivity of the interaction. Larger *γ* means that the predation probability decreases faster with increasing phenotypic mismatch between the prey and the predator. Whether or not predation does occur at any given encounter is then determined by a Bernoulli draw with probability *I*(*x, y*). In case of successful predation, a variable *d* of the individual prey is set to 1, tagging it for removal by death (thus preventing it from further interacting with predators), and a variable *c* of the individual predator is incremented by 1, thus increasing the number of prey it has consumed. After all encounters and predation events are processed, the predatory component of fitness is updated for each individual prey (*V*_*x*_) and predator (*V*_*y*_) as

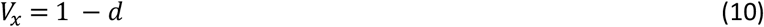

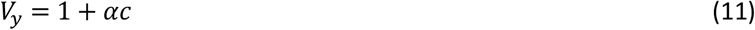

where the reproductive pay-off *α* quantifies how much additional fitness (number of offspring) is gained by the predator for each additional prey consumed.

#### Demography

We wish to compare purely (co)evolutionary models where the population sizes of interactors are unaffected by their evolution, with eco-evolutionary models where evolution and demography interact (67), possibly leading to eco-evolutionary feedbacks (68). We thus develop both a coevolutionary model with constant population sizes, and another with population size dynamics. In both cases, we assume discrete non-overlapping generations, wherein hermaphroditic diploid individuals reproduce sexually and with random mating, with self-fertilization permitted (see Table S1 for the parameter values associated to each demographic model).

We first simulate Wright-Fisher populations, where the numbers of prey and predators are fixed at *N* and *P* in all generations. This amounts to assuming that the total number of offspring produced is always larger than the number of parents, and that strong ceiling-type density regulation takes place in both species. Under these assumptions, the next generation of prey is generated by randomly drawing two parents for each of *N* offspring individuals, using the fitness *W*_*x*_ of each parent as weight for the sampling probabilities (and similarly for predators using *W*_*y*_).

In the eco-evolutionary model, the life cycle is more explicit, and fitness directly influences population growth. For the prey, *W*_*x*_ (as specified above) determines the probability of survival to reproduction. Following this first episode of stabilizing selection, each surviving (hermaphroditic) individual reproduces as a female, by drawing a random individual from the population as male. We model density-dependent fecundity using the Beverton-Holt model (a discrete-time version of logistic growth), wherein the expected number of offspring per female is 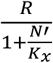 with *R* the intrinsic multiplicative growth rate, *N′* the number of prey adults that survived to reproduce, and *K*_*x*_ the carrying capacity (expected number of individuals, neglecting mortality before reproduction). Demographic stochasticity (between-individual randomness in reproductive output) is introduced by drawing the number of offspring for each couple from a Poisson distribution, with the expected number of offspring as parameter. For predators, the main difference is that only the stabilizing selection term *U*_*y*_ determines the survival probability, while *V*_*y*_ determines the predation success, which influences the expected fecundity. As for prey, the realized fecundity is also influenced by density-dependent regulation and demographic stochasticity. Specifically, the number of offspring per reproducing female is drawn from a Poisson distribution with parameter 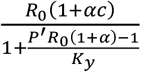, where *R*_0_ is the baseline intrinsic multiplicative growth rate, *P′* the number of predator adults that survived to reproduce, and *K*_*y*_ is a parameter that tunes the strength of density regulation. Note that the equilibrium population size of predators depends both on *K*_*y*_ and on the intrinsic growth rate *R*_*O*_*(1 + αc)*, but the latter is not constant unlike for prey, as it depends on the number of consumed prey. Hence *K*_*y*_ corresponds to the equilibrium population size of a putative population where each predator would catch one prey on average, leading to an intrinsic growth rate *R*_*O*_*(1 + α)*.

#### Environmental fluctuations

We incorporated environmental fluctuations that shift the optimal phenotype for the prey, and possibly also for the predator. We first considered periodic fluctuations, modeled as a sine wave,

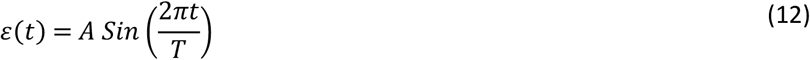

where *ε(t)* is the environmental value at time step *t*, and *A* and *T* are the amplitude and period of fluctuations, respectively. This collapses to a constant environment at *ε(t) = 0* when setting *A* to 0 (see Table S1 for other explored values for *A and T*). We also considered stationary stochastic fluctuations, in the form of a first-order autoregressive process with variance 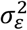 and autocorrelation *ρ* over one time step. This was modeled by drawing the first environment *ε*(0) from a normal distribution *N*(0, *σ*_*E*_), and all following environments (for *t* > *0*) as

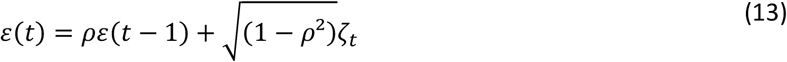

where the *ζ*_*t*_ are drawn from independent normal distributions *N(0, σ*_*E*_). The prey’s and predator’s optimum phenotypes without predation are then respectively set as

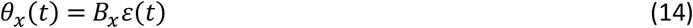

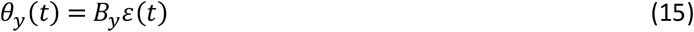

where *B*_*x*_ and *B*_*x*_ are the slopes of changes in the optimum phenotypes for prey and predators with the environment (described as environmental sensitivity of selection in ref. (69)). For instance, *B*_*y*_ *= 0* means that the optimum phenotype of the predator is unaffected by environmental fluctuations and remains constant, while *B*_*x*_ *= B*_*y*_ means that the optimum phenotypes of the predator and the prey are affected in the same way by the fluctuating environment. The parameter values used for *B*_*x*_, *B*_*y*_, *σ*_*E*_ and *ρ* appear in Table S1.

#### Scenarios

We wanted to understand how adaptation to a moving optimum in prey is affected by the presence of predators that do or do not adapt to the changing environment. We therefore contrasted scenarios that differed with respect to predator presence and evolution, summarized in Table 1. In the ‘no-predator’ scenario (NP), the prey were simulated in isolation, without predators (*P = 0* throughout). In the ‘non-selective predator’ scenario (NSP), the predator hunted non-selectively, that is, the probability of predation was determined independently of the phenotypic difference with the prey (*I(x, y) = M* for all *x* and *y*). In the ‘non-evolving predator’ (NEP) scenario, predation was conditioned on phenotypic similarity, but the predators’ trait remained constant at a value corresponding to the average optimum of the prey (*S*_*y*_ *= B*_*y*_ *= 0*). In the ‘evolving predator with constant optimum’ (EPC) scenario, the selective predator could evolve, but its optimum phenotype was not affected by the fluctuating environment, instead remaining constant at a value corresponding the mean optimum for the prey (*B*_*y*_ *= 0*). Lastly, in the ‘evolving predator with fluctuating optimum’ (EPF) scenario, the predator evolved and its optimum phenotype changed in the same way as that of the prey (*B*_*y*_ *= B*_*x*_ *= 1*).

#### Simulations and data analysis

A first set of simulations was conducted under a Wright–Fisher model and with a constant environment (i.e., constant *θ*_*x*_ and *θ*_*y*_), for various parameter values (*γ, S*_*x*_, *S*_*y*_, *M*, and *α*; detailed in Table S1). In all cases, *θ*_*y*_ was fixed at 0, while *θ*_*x*_ varied from 0 to 40 by increments of 1, allowing exploration of a range of differences between optimum phenotypes in the prey and the predator. Each simulation included a burn-in period of 500 generations. The mean trait values of the prey and predator were then recorded at each generation, and averaged over the final 500 generations. Three replicates were run for each condition.

A second set of simulations was conducted with a periodically fluctuating environment, for all 5 scenarios introduced above, still under a Wright–Fisher model. We used a fixed set of parameter values for the strengths of stabilizing selection and predatory interactions (*γ = 0*.*01, S*_*x*_ *= 0*.*01, S*_*y*_ *= 0*.*01, M = 0*.*6* and *α = 1*), and varied the amplitude *A* of oscillations from 0 to 40 by increments of 1, and their period *T* as detailed in Table S1. To quantify how well prey tracked their optimum in a changing environment, we computed the amplitude of cycles in deviations of the mean phenotype from the optimum. The difference between the mean phenotype of prey and their optimum phenotype was recorded in every generation for 100*T* generations, following a burn-in time of 5*T generations*. We then averaged this deviation from the optimum over the 100 periods for each generation in the cycle (that is, averaging generations *t, t + T, t + 2T, …* , *t + 100T*, for all *t* between 1 and *T*), to remove the noise caused by random genetic drift. We used a single simulation run, relying on the ergodicity afforded by the large number of periods. The amplitude of this average cycle in deviations from the optimum in prey was then estimated as 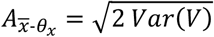, where *V* is the vector of averages over all generations in a period. This formula uses the fact that the variance of a sine wave with mean 0 (as occurs for deviations from the optimum) is 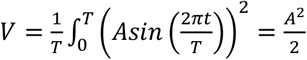. Therefore, this approximation is expected to be less accurate under conditions where cycles in deviations from the optimum do not conform to a sine wave, for instance when changes in the strength of predatory selection lead to strong non-linearity in evolutionary responses (Fig. S6). We also computed the average genetic variance over these 100 periods. For the NP and NSP scenarios, we additionally computed the (inbreeding) effective population size of prey in every generation as

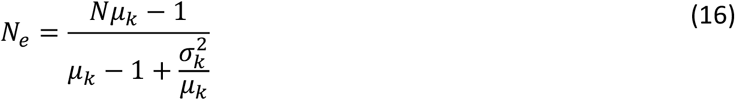

with *N* the population size, *μ*_*k*_ the mean offspring number (which is always 2 under constant population size) and 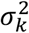 the variance of offspring number (eq. 7.6.4.1 in ref. (70), eq. 7 in ref. (71)). The long-term effective population size was then computed as the harmonic mean of *N*_*e*_ across the 100 periods (70, 71).

A third set of simulations was conducted with a stochastically fluctuating environment (first-order autoregressive process), for all 5 scenarios introduced above, under a Wright–Fisher model (Tables 1 and S1). We used the same set of parameter values as in the periodically fluctuating case, and varied the standard deviation of the environment *σ*_*E*_ from 0 to 40 by increments of 1, and the auto-correlation of the environment *ρ*. For consistency with the periodic case and previous literature (31), we parameterized autocorrelation in terms of a characteristic time *T*, which we define as the time after which autocorrelation becomes lower than 1%, such that *ρ*^*T*^ *= 0*.*01*, where the autocorrelation over one generation is 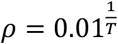. To quantify how well prey tracked their optimum in a stochastically fluctuating environment, we recorded the deviation between the mean phenotype and the phenotypic optimum in every generation over *500T* generations (following a burn-in period of *5T* generations), and computed their standard deviation. We averaged over longer times than in the periodic case because environmental stochasticity introduces an additional component of noise in evolution (beyond random genetic drift and the randomness of mutation events (72)).

Similar sets of simulations were run for an eco-evolutionary model with explicit demography (non Wright-Fisher in SLiM). The initial population sizes were set to 10000 for the prey and predators, and we used the same selection parameters as in Wright–Fisher simulations, with additional parameters detailed in Table S1. Under a constant environment, unlike in the Wright–Fisher model, prey optimum *θ*_*x*_ was progressively (i.e., every 1000 generations) increased by steps of 1 from 0 to 40. This procedure allowed reducing the extinction risk that could be induced by a drastic environmental shift from 0 to a very different optimum, while still allowing the system to reach its evolutionary equilibrium at the end of each step. The mean phenotypes and population sizes after reproduction were then recorded and averaged for prey and predators over 500 generations, after a burn-in period of 500 generations. Three replicate simulations were run for each condition.

Periodic environmental cycles for all 5 scenarios were also investigated in the eco-evolutionary model (Tables 1 and S1), using the same fixed set of parameter values for selection (*γ = 0*.*01, S*_x_ *= 0*.*01, S*_*y*_ *= 0*.*01, M = 0*.*6* and *α = 1*), the period *T* as detailed in Table S1, and progressively increasing the amplitude *A* by steps of 1 from 0 to 40. The population sizes, mean phenotypes, deviations from optimum (i.e., mean phenotype minus optimum value), and genetic variance of prey, were recorded before selection in every generation, over 100 periods of a single simulation run, after a burn-in of five period, for each assayed period and amplitude. The amplitude of the deviation from optimum in prey was computed as detailed above for the Wright–Fisher model.

Graphic displays were performed using R software v 4.3.2.

## Supporting information

Supplementary Mathematica notebook

## Codes and data availability

Codes used to generate data and figures are available at: https://gitlab.com/Louguyot/coevolving_predators_and_prey_adaptation_guyot_2025

## Acknowledgements

We thank the GEE team at Ecologie, Société, Evolutibon for providing access to their computing resources, and two anonymous reviewers for useful feedback.

## Supplementary Appendix

**Analytical approximations**

To help interpret the simulation results and relate them to earlier findings from theory not based on individual-based processes, we relied on a simplified version of the model that only tracks population-level variables, such as the mean and variance of traits, and population sizes of interactors. This population-based model was derived by combining the theory of phenotype-matching coevolution (34–36) with models of adaptation to a fluctuating optimum (31, 32, 51). However, a key difference with earlier coevolutionary theory is that we explicitly let selection emerge from individual processes of encounters and interactions, which led to qualitatively different results. Some of the detail of the derivations below can be found in the Supplementary Mathematica notebook.

### Selection components

The first step involves deriving the expected predatory fitness of phenotype *x* in prey, or *y* in predators, from events occurring at the individual level. For the prey, summing over all possible numbers of encounters with predators weighted by their Poisson distributed probability, using eqs. (9) and (10) in the main text the expected survival probability of individuals with phenotypes *x* conditional on the phenotypes *y* of the predator is

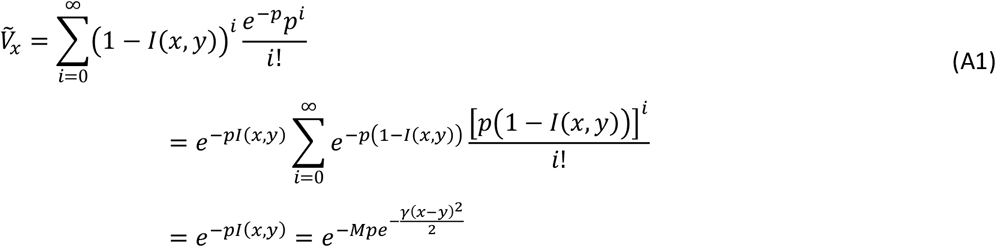

where the tilde denotes an average over prey individuals with phenotype *x*, and we have used the fact that the second line includes a sum of terms from a Poisson distribution (with parameter *p*(1 ™ *I*(*x, y*))), which equals 1 by definition of a probability distribution.

Assuming that the phenotypes of predators and prey are normally distributed with small variances, a similar equation holds approximately for the mean predatory fitness of prey in terms of their mean phenotype 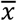 (and conditional on the mean phenotype of predators 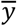), which we denote as 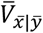. The selection gradient on the prey’s trait caused by interactions with the selective predator is obtained by taking the derivative of the logarithm of this mean fitness component relative to 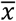, leading to

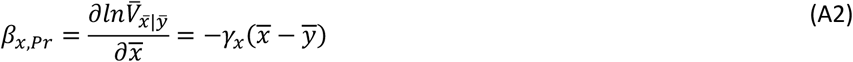

where the index ‘Pr’ denotes the predatory component of selection, and the intensity of predatory selection on prey (conditional on phenotypic mismatch) is

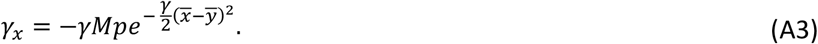

Equation (A2) mirrors similar ones from previous trait-matching models of coevolution defined at the population level (34–36), except for a factor 2 caused by a slightly different parameterization (as some of these studies defined the Gaussian interaction function without 1/2 in the exponential). Consistent with these earlier models, we have *γ*_*x*_ *< 0* regardless of phenotypic mismatch (from eq. A3), which indicates that the prey is always selected to evolve away from the phenotype of the predator, thus reducing the intensity of predation. However there is an important difference with population-based models of coevolution. Whereas these earlier models assumed a constant intensity of predatory selection *γ*_*x*_, in our model that starts from first principles at the individual level, *γ*_*x*_ instead depends on the mismatch 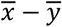 between the predator and prey mean phenotypes, through the term 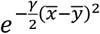 in eq. (A3).

For predators, summing over all possible numbers of consumed prey, assumed to be Poisson distributed with rate given prey density 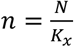 (with *K*_*x*_ the carrying capacity of prey) weighted by predatory success, using eqs. (9) and (11) in the main text the expected number of offspring of individuals with phenotype *y* is

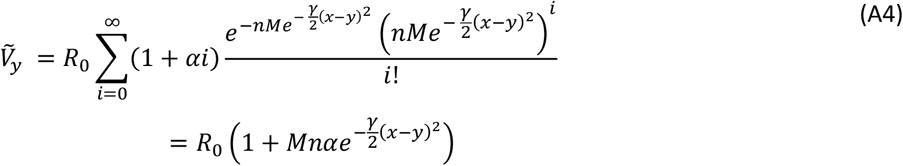

where the tilde denotes an average over predator individuals with phenotype *y*, and we have used the expectation of a Poisson distribution. Again assuming normal phenotype distributions with small genetic variances, the expected mean predatory fitness of predators 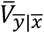 is proportional to the right member of eq. (A4), with *x* and *y* replaced by their population means 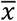 and 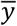. Taking the derivative of the logarithm for this expression, the selection gradient on predators caused by selective predation on the prey is.

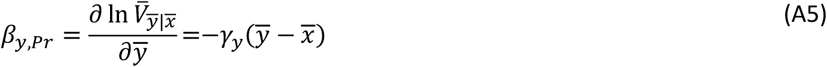

where the intensity of predatory selection on the predator (conditional on phenotypic mismatch) is

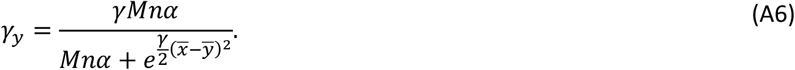

As assumed in earlier less mechanistic models (34–36), *γ*_*y*_ > *0* in all cases, such that the predator is always selected to track the phenotype of its prey (from eq. A5), thereby increasing the intensity of predation. However like *γ*_*x*_ , and contrary to the assumptions from most individually implicit models, *γ*_*y*_ here depends on the mean phenotypic mismatch 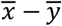.

For stabilizing selection, also assuming normally distributed traits with small phenotypic variance relative to the width 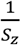 of the fitness peak, the mean fitness component caused by adaptation to the abiotic environment 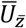 is proportional to 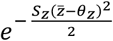 (from eq. 8 in the main text and ref (44)). Taking the derivative of the logarithm for this expression, the associated selection gradient is

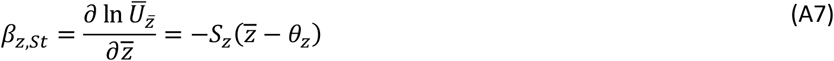

where the index ‘St’ denotes the stabilizing component of selection, *z = x* for prey and *z = y* for predators. Equation (A7) shows that adaptation to the abiotic environment leads to a component of directional selection that brings the mean phenotype towards its optimum, proportionally to how much this mean phenotype deviates from the optimum (49).

#### Response to selection in a given generation

Because the two components of selection imposed by the biotic and abiotic environment affect fitness multiplicatively, due to properties of logarithms their combined effects on directional selection are obtained by summing their selection gradients (73), such that the expected response to selection is well approximated by

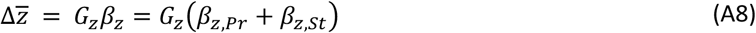

where *β*_*z*_ is the total selection gradient, *G*_*z*_ is the additive genetic variance, and again *z = x* for prey and *z = y* for predators (34, 35). Replacing with the expressions for selection gradients in eqs. (A2) and (A7), the total selection gradient in prey becomes, after some rearrangement,

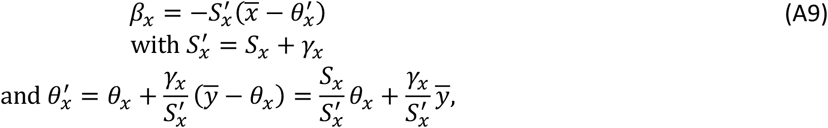

consistent with earlier results from the literature (34–36). An analogous result holds for predators, simply inverting the *x* and *y* subscripts. Equation (A9) shows that when selective predation is combined with selection for an optimum phenotype exerted by the non-predatory environment, as long as 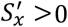 the evolutionary dynamics are similar to those without predation (compare eq. A9 with eq. A7), but with a modified effective strength of stabilizing selection 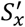, and a displaced effective optimum 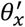. Since *γ*_*x*_ is negative for prey (from eq. A3), the effective strength of stabilizing selection in prey is reduced by selective predation (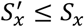, broader fitness peak than without predation). Very strong predatory selection, such that |*γ*_*x*_| > *S*_*x*_, could even turn selection from stabilizing to disruptive (fitness valley) as *γ*_*x*_ is negative, possibly leading to runaway dynamics (34), but we do not investigate this scenario here because it does not lead to adaptive tracking of a moving optimum.

Because *γ*_*x*_ is negative for prey (from eq. A3), the effective optimum for prey 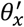 is shifted *away* from the predator’s mean phenotype 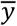 (evolutionary escape), to an extent that depends on how much 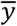 deviates from the prey’s optimum without predation *θ*_*x*_, and on the relative strengths of predatory over stabilizing selection 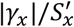. Conversely for predators *γ*_*y*_ is positive (from eq. A6), so the effective strength of stabilizing selection is always increased by selective predation (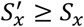, narrower fitness peak than without predation). The effective optimum for predators is shifted *towards* the prey’s mean phenotype 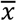(evolutionary chase), to an extent that depends on how 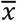 deviates from the predators optimum *θ*_*y*_ and the ratio of selection strengths 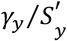.

A joint equilibrium for the mean phenotypes in predators and prey occurs for 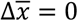 and 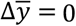. Combining eqs. (A8) and (A9), this condition leads to the system of equations

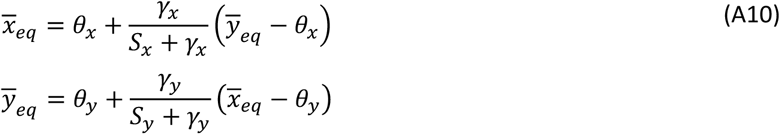

Inserting the expression for 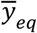 into the first equation (and reciprocally for 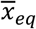 into the first equation) leads to

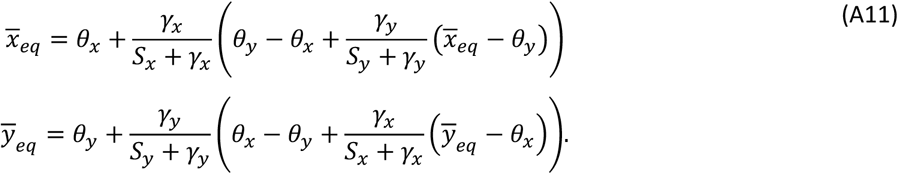

This can be rearranged as

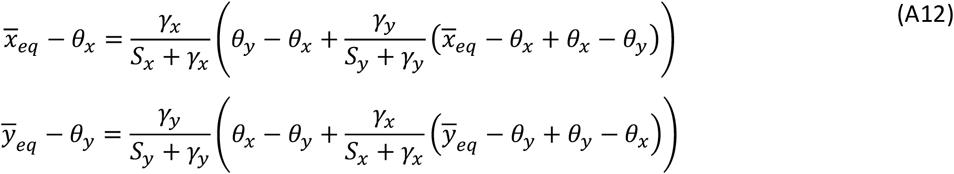

such that

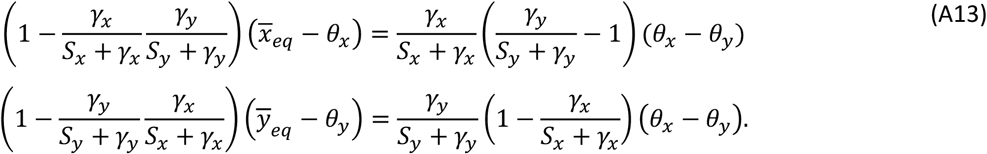

Further simplifying we get

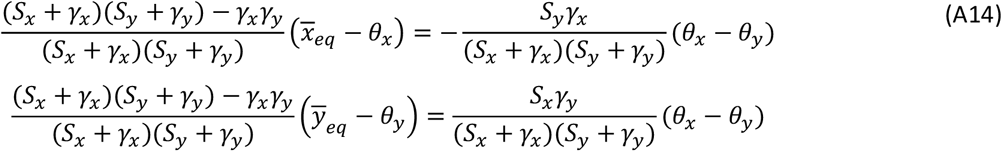

then

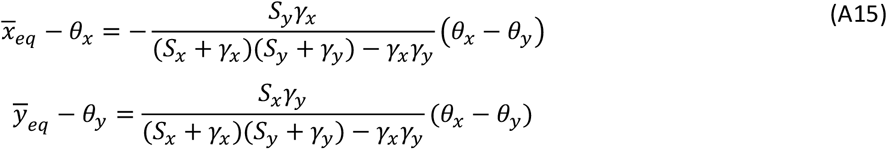

and finally.

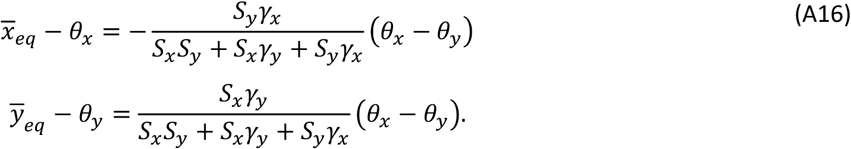

Dividing the numerator and denominator by *S*_*x*_*S*_*y*_ for both equations then leads to

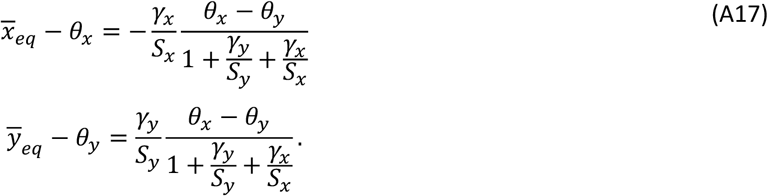

From this we get that

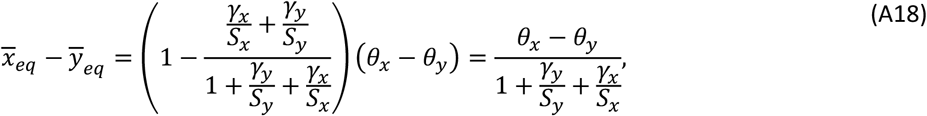

which leads to eqs. (1) and (2) in the main text.

The equilibria in eqs. (A17-A18) are not fully explicit, because the strengths of predatory selection *γ*_*x*_ and *γ*_*y*_ depend on the phenotypes of prey and predators (from eqs. A3 and A6), leading to feedbacks that preclude predicting the coevolutionary equilibria from basic parameters of the model. However, when predatory selection is sufficiently weak relative to stabilizing selection in both species, that is when 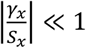 and 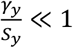, we get from eq. (A18) that 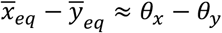. We may thus replace 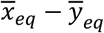 by *θ*_*x*_ − *θ*_*y*_ in the expressions for *γ*_*x*_ and *γ*_*y*_ in eqs. (A3) and (A6), leading to eq. (5) and (6) in the main text. When this approximation holds, the coevolutionary equilibrium for the mean phenotypes of prey and predators can be expressed in terms of the primary parameters of the model. Under these same assumptions that 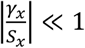 and 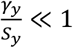, the deviations from the optimum in both the prey and the predator become proportional to 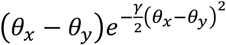 for large |*θ*_*x*_ − *θ*_*y*_| (inserting eq. 5 and 6 into eq. (A18) with the denominator approximated as 1, and taking the limit for large |*θ*_*x*_ − *θ*_*y*_|). This expression tends to 0 as |*θ*_*x*_ − *θ*_*y*_| tends to infinity (using L’Hôpital’s rule), such that the deviation from the optimum in both predators and prey vanishes as the divergence between their optima becomes larger.

#### Evolutionary response to a changing environment

In a periodic environment, inserting 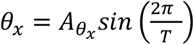 (with the amplitude of the optimum being 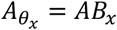, from eqs. 12 and 14 in the main text) in eq. (A8-A9), we get

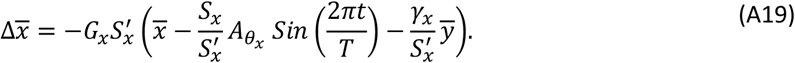

Let us first consider the case where the predator’s mean phenotype 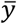 remains constant at the average optimum of the prey (set to 0), as assumed in scenario NEP (and also approximating the scenario EPC when the predator barely tracks the prey’s mean phenotype). The per-generation change in the mean phenotype of the prey then becomes

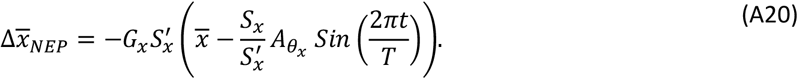

Equation (A20) shows that in this case, prey effectively track an optimum that undergoes cycles with larger amplitude than that of the optimum (by a factor 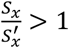), but do so with a reduced effective strength of selection 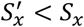, as compared to the case without a selective predator (obtained by setting *γ*_*x*_ *= 0* such that 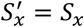). Similarly in a stochastic environment, the prey effectively tracks fluctuations with a larger standard deviation, by a factor 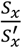, with a reduced selection strength 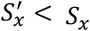.

In the opposite case where the predator tracks the same optimum as the prey (scenario EPF), we need to account for changes in the mean phenotype of predators 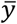 when analyzing selection and evolution of prey. Previous theory without species interactions (31, 32, 51) suggests that 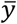 should eventually undergo cycles with the same period as the optimum, but amplitude reduced by a factor 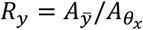, and phase lagging by a proportion *φ*_*y*_ of a cycle. Assuming that this also holds (at least approximately) with species interactions – consistent with Fig. S5, and with the fact that tracking of the prey’s mean phenotype should mostly “pull” the predator’s mean phenotype closer to the optimum – we can substitute 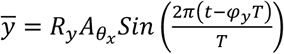 into eq. (A19), yielding

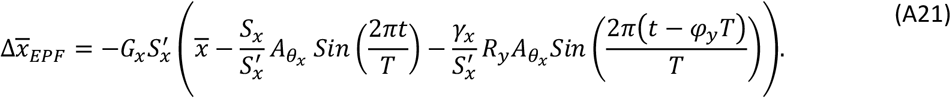

A linear combination of sine waves with the same period can be rearranged as a single sine wave with the same period as both, but with a different amplitude and shifted phase, such that we have

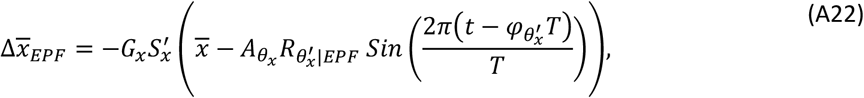

where 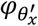 is the phase shift of the prey’s effective optimum relative to the true optimum (expressed as a proportion of a full cycle), and 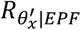 is the relative amplitude of the effective optimum scaled that of the true optimum. To obtain the latter, we derive the expectation of the squared linear combination of sines over a cycle, multiply it by two, and take the square root (see the Supplementary Mathematica notebook for more detail), leading to

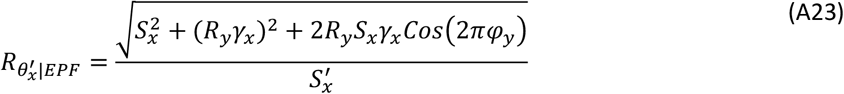

More insights can be gained by further analyzing cycles in the predator’s mean phenotype 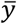. Earlier theory without coevolution (31, 32, 51) has shown that the asymptotic cycle in the mean phenotype has smaller amplitude than cycles in the optimum (*R*_*y*_ ≤ *1*), and a phase lag between 0 (under efficient adaptive tracking of the optimum) and a quarter period (under poor adaptive tracking). Since adaptive tracking in the predator is expected to be improved by the selective pull exerted by the mean phenotype of prey, we also expect that 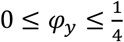 under EPF. Taking the limit under efficient adaptive tracking by the predator (*φ*_*y*_ → *0*) leads to

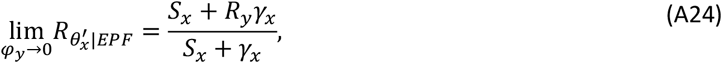

where we have used the fact that *S*_*x*_ *+ R*_*y*_*γ*_*x*_ > *0* under our assumption that |*S*_*x*_| > |*γ*_*x*_| (required so that the net selection on 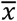 is stabilizing, and thus compatible with tracking a moving optimum). Equation (A24) shows that in the limit of very efficient tracking by the predator, the prey effectively tracks an optimum with larger amplitude than its true optimum (since *γ*_*x*_ *< 0* and *R*_*y*_ ≤ *1*). The limit under poor adaptive tracking by the predator 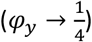 is

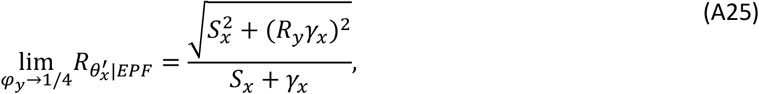

which may be larger or smaller than 1 depending on *R*_*y*_.

The expressions in eqs (A23-A25) are for the amplitude of cycles in the effective optimum for prey 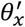, but ultimately maladaptation of prey with respect to the non-predatory environment (as measured by e.g. tolerance curves in the laboratory) depends on how much their mean phenotype deviates from their true optimum *θ*_*x*_. The mean phenotype of prey tracks the effective optimum 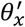 (influenced by selective predation) through adaptive evolution, and this determines how this mean phenotype deviates from the predation-independent optimum *θ*_*x*_. We can obtain further analytical insights by assuming a constant genetic variance *G*_*x*_ as in previous theory without coevolution (31, 32, 51), and also assuming that *γ*_*x*_ remains constant throughout the cycle (that is, assuming eq. (3) in the main text always holds), such that 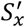 is also constant. We can then extend results from earlier theory on adaptation to a cycling optimum without predators (31, 32, 51) to derive analytical approximations for the amplitude of cycles in deviations of the mean prey phenotype from its optimum *θ*_*x*_, which determine the average fitness over a cycle (31, 32, 51). The details of the derivation can be found in earlier theoretical literature on adaptation to a cycling environment without a predator (31, 32, 51), and are here transposed to the case with a predator, where the prey tracks an effective optimum 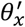. In brief, turning eqs. (A19-A22) into differential equations (that is, replacing 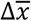 by 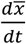, which is a good approximation under weak selection as assumed throughout our study), solving for 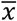, and taking the asymptotic limit for large times, yields a linear combination of sine and cosine waves, with the same period *T* as the optimum. From trigonometric identities, this can be rearranged as a single sine wave,

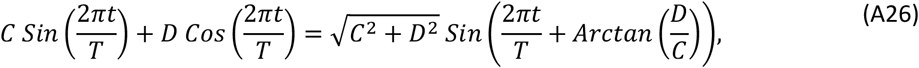

with the same period *T*, but phase shifted by 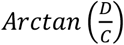, and amplitude 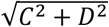. Finally, since the difference between the sine waves for the prey’s mean phenotype 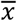 and optimum phenotype *θ*_*x*_ can also be rearranged as a single sine wave, its amplitude can be derived as detailed before eq. (A23), by taking the square root of twice the mean square of this sine function over a cycle. The detailed derivations in each specific scenario appear in the Supplementary Mathematica notebook.

In the scenario NEP where the predator does not evolve, we find that the amplitude of deviations from the optimum scaled to the amplitude of the optimum itself 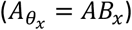 equilibrates in the long run to

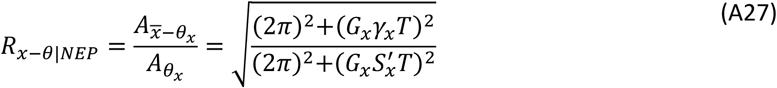

The ratio of the amplitude of deviations from the optimum with a constant predator (NEP) to that without a predator (NP) is

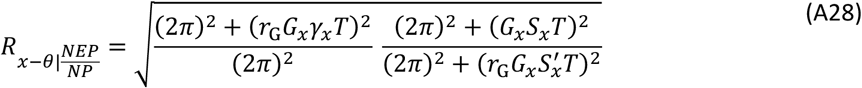

where *r*_*G*_ is the ratio of genetic variances with versus without the predator. Inspection of eq. (A28) indicates that the first ratio is always larger than 1 (since *(r*_*G*_*G*_*x*_*γ*_*x*_ *T)*^*2*^ ≥ *0*), while in the second ratio we have 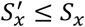 (recalling that *γ* ≤ *0* for the prey) but *r* ≤ *1* or *r* ≥ *1* depending on the conditions (Fig. 3C and S7B). Further insights can again be obtained by focusing on limit cases. Under parameter values leading to weak responses to selection in a rapidly fluctuating environment (such that *(r*_*G*_*G*_*x*_*γ*_*x*_*T)*^*2*^ ≪ *(2π)*^*2*^, *(G*_*x*_*S*_*x*_*T)*^*2*^ ≪ *(2π)*^*2*^, and 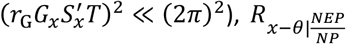 tends to 1. In the reverse situation of substantial responses to selection in a slowly fluctuating environment 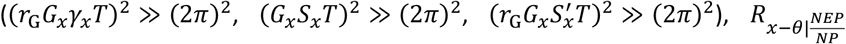 instead tends to 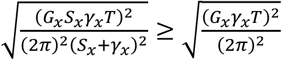 (where we have used the fact that 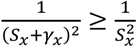, which is larger than 1 under these assumptions, and increases with increasing (co)evolutionary potential. Hence a non-evolving predator with phenotype fixed at the mean optimum of the prey is expected to amplify deviations from the optimum in the prey under conditions where responses to selection are efficient.

The case with a co-adapting predator (EPF) is more complex. Assuming as we did above that the predator’s mean phenotype undergoes a sine wave with reduced amplitude and lagged phase (consistent with Fig. S5), the ratio of amplitudes of deviations from the optimum in this scenario relative to the case without a predator (NP) is

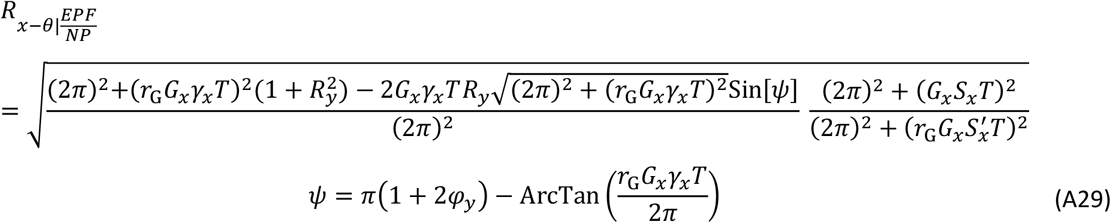

Comparing with the expression above for NEP (eq. A28), the main difference is that the term *(r*_*G*_*G*_*x*_*γ*_*x*_*T)*^*2*^ in the first fraction is now replaced by a more complex expression that depends on the parameters of fluctuations in the mean phenotype of the predator (*φ*_*y*_ and *R*_*y*_). Setting *R*_*y*_ to 0 (no fluctuations in the predator’s mean phenotype) recovers eq. (A28), as it should. As indicated below eq. (A23), we expect from earlier theory without coevolution (31, 32, 51) that *R*_*y*_ ≤ *1* and 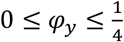, which should also hold in the EPF scenario, since adaptive tracking in the predator is improved by the selective pull from the prey. Substituting these boundary conditions for *φ*_*y*_ in the expression for *ψ* in eq. (A29) leads after some algebra to

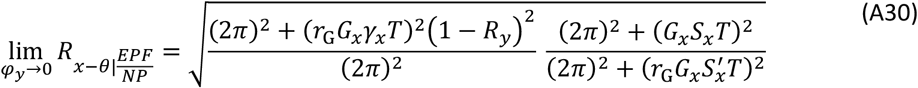

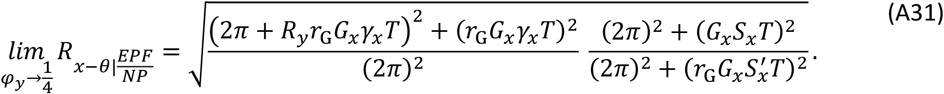

The limit of perfect tracking in the predator (*φ*_*y*_ → *0*) is the most different from scenario NEP. It leads to a strikingly similar expression as in this scenario (comparing eq. A30 with eq. A28), except for a multiplication by *(1* − *R*_*y*_*)2* in the first ratio in eq. (A30). Since *R*_*y*_ can be of order 1 (but still below 1) under efficient tracking, this means that maladaptation in the prey can be very much reduced when the predator closely tracks the optimum, as compared to when the predator’s mean phenotype remains constant. Whether maladaptation is also reduced relative to the case with no predator depends on other parameters values such as the ratio of genetic variances, for which no analytical solutions exist.

## Supplementary Tables and Figures

**Table S1:** Parameter values explored in the simulations. Predatory parameters *S*_*y*_, *B*_*y*_ and *P* are set to 0 in specific scenarios of changing environment (detailed in Table 1), and stars denote parameter values only explored in supplementary figures.

| Parameters | Explored values |  |  |  |  |
| --- | --- | --- | --- | --- | --- |
| Model | Wright-Fisher |  |  | Eco-evolutionary dynamics |  |
| Environment | Constant | Periodic | Stochastic | Constant | Periodic |
| Mutation rate per locus ($\mu$) | $10^{-5}$ | | | | |
| Number of loci ($L$) | 1000 | | | | |
| Recombination rate between adjacent loci ($r$) | $10^{-3}$ | | | | |
| Prey population size ($N$) | 10000 | | | - | |
| Predator population size ($P$) | 10000 | | | - | |
| Carrying capacity for prey and predator ($K_x, K_y$) | 10000 | | | | |
| Strength of stabilizing selection for prey and predator ($S_x, S_y$) | (0.01, 0.01) / (0.1, 0.1)* | (0.01, 0.01) | | (0.01, 0.01) / (0.1, 0.1)* | (0.01, 0.01) |
| Selectivity of the interaction ($\gamma$) | 0.01 / 0.1* | 0.01 | | 0.01 / 0.1* | 0.01 |
| Prey optimum ($\theta_x$) | 0 to 40 | - | | 0 to 40 | - |
| Predator optimum ($\theta_y$) | 0 | - | | 0 | - |
| Maximum predation intensity ($M$) | 0.2 / 0.6 / 0.8 | 0.6 | | 0.2 / 0.6 / 0.8 | 0.6 |
| Reproductive pay off of predation ($\alpha$) | 0.25 / 1 / 4 | 1 | | 0.25 / 1 / 4 | 1 |
| Intrinsic multiplicative growth rate of prey ($R$) | - | | | 3 | |
| Intrinsic multiplicative growth rate of predator ($R_0$) | - | | | 1 | |
| Amplitude of environmental cycles<br>($A$) | 0 | 0 to 40 | - | 0 | 0 to 40 |
| Period of environmental cycles /<br>Characteristic time of stochastic<br>fluctuations ($T$) | - | 50 / 100 / 200 / 400 | | - | 50 / 100 /<br>200 / 400 |
| Environmental standard deviation<br>($\sigma_\varepsilon$) | - | - | 0 to 40 | - | - |
| Environmental sensitivity of<br>selection in prey and predators<br>($B_x, B_y$) | - | 1 | | - | 1 |

**Figure S1:**
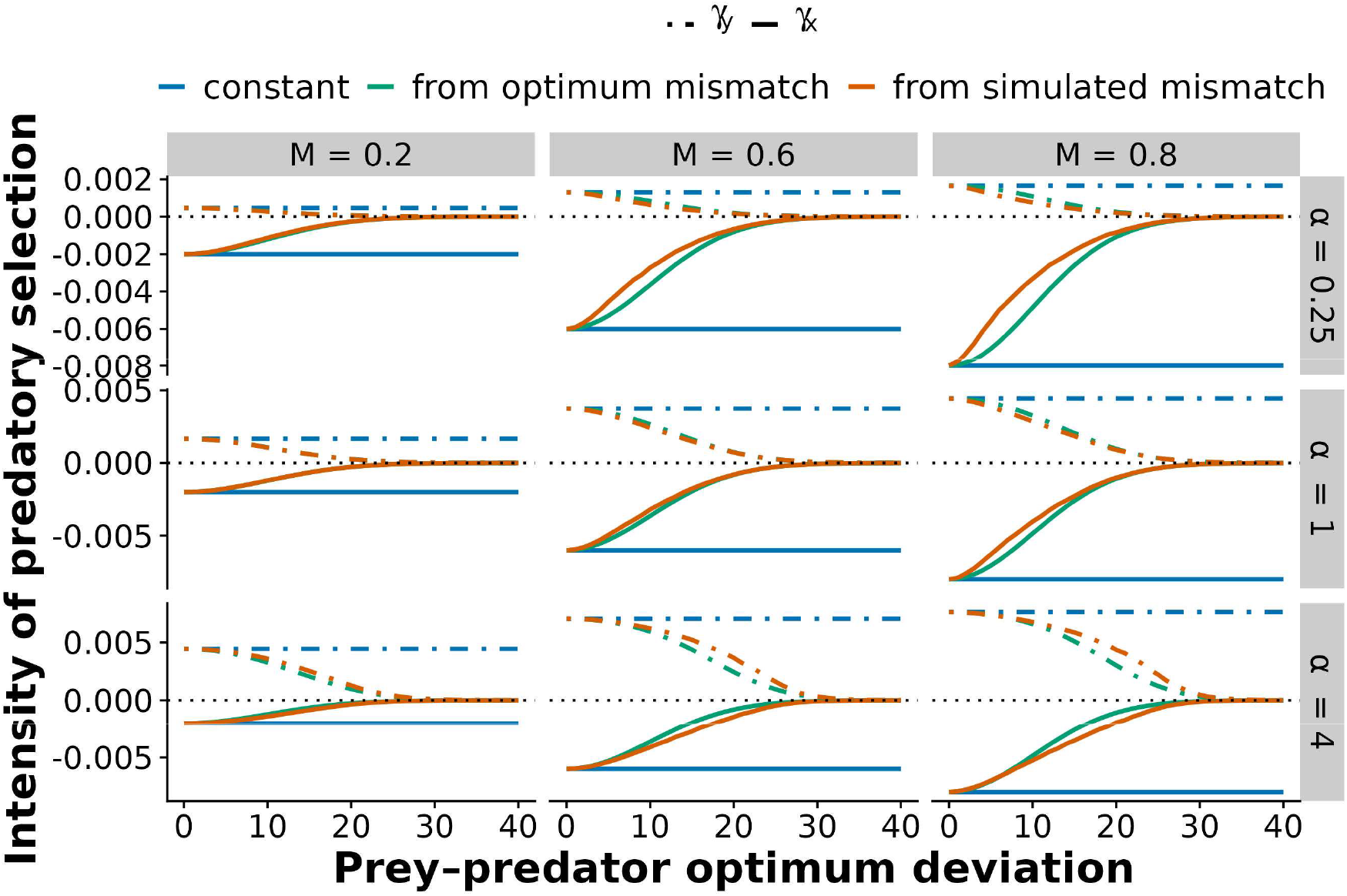
Intensity of predatory selection in a constant environment with constant population sizes. The predicted intensity of predatory selection in prey (*γ*_*x*_, continuous lines) and in predators (*γ*_*y*_, dashed lines) is plotted against the difference between the optimum phenotypes in prey and predators, using various approximations: eqs. (3, 4) (blue), eqs. (5, 6) (green), or eqs. (A3, A6) using the mean mismatch from three replicate simulations (orange). The parameters used are as in the main text: *γ = 0*.*01; N = 10000; P = 10000; S*_*x*_ *= 0*.*01; S*_*y*_ *= 0*.*01* and *M* and *α* as indicated. The black dotted line indicates 0.

**Figure S2:**
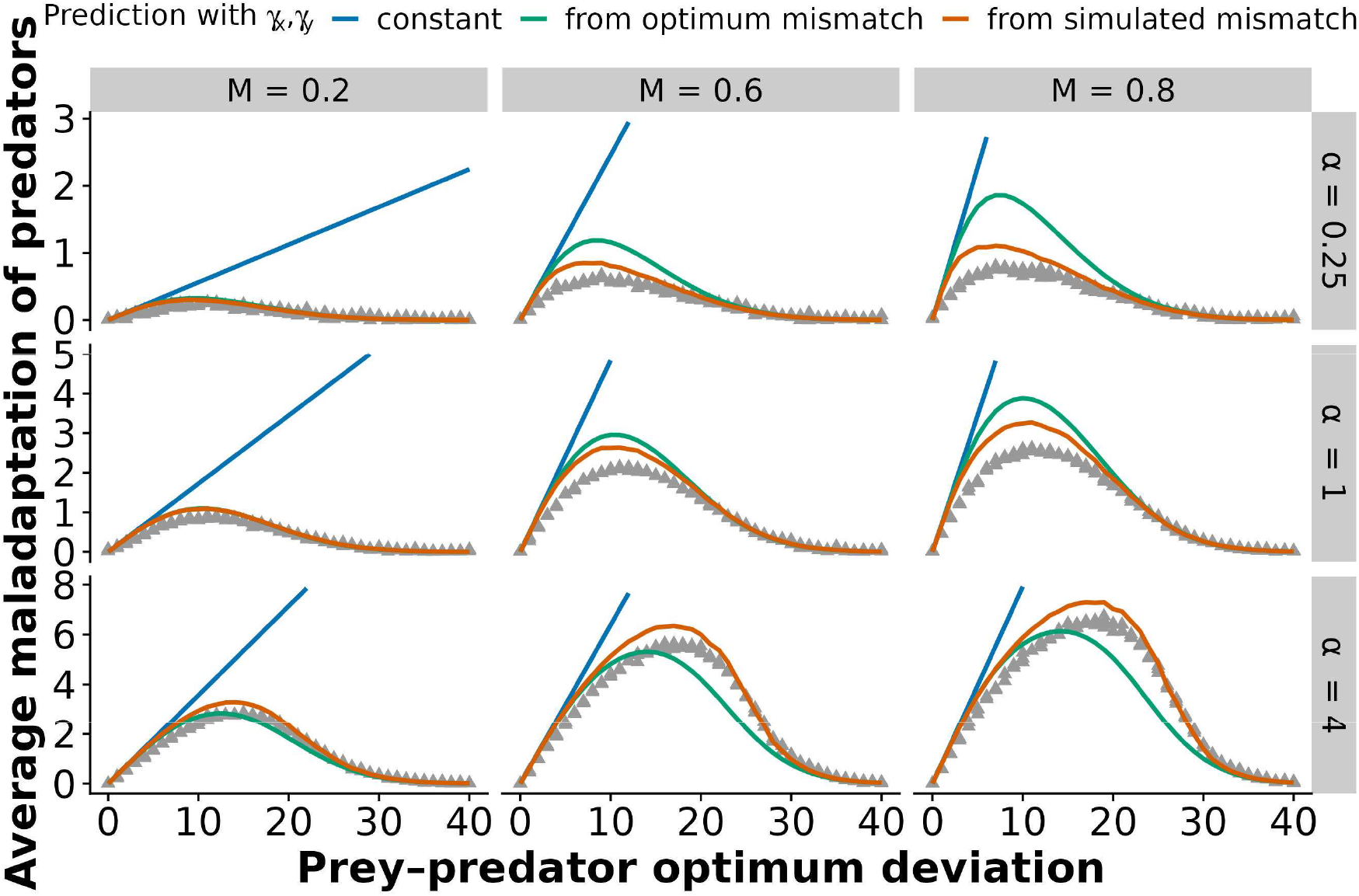
Maladaptation in predators in a constant environment with constant population sizes. The phenotypic deviation from optimum in predators is plotted against the difference between the optimum phenotypes in prey and predators. Gray triangles show three replicate simulations of Wright– Fisher populations, averaged over 500 generations after a burn-in period of 500 generations, across a range of optimum differences (from 0 to 40). Full lines show predictions from eqs. (1, 2), using different approximations for *γ*_*x*_ and *γ*_*y*_: eqs. (3, 4) (blue), eqs. (5, 6) (green), or eqs. (A3, A6) using the mean mismatch from three replicate simulations (orange). Simulations were performed with *γ = 0*.*01; N = 10000; P = 10000; S*_*x*_ *= 0*.*01; S*_*y*_ *= 0*.*01*, and *M* and *α* as indicated in each column and row.

**Figure S3:**
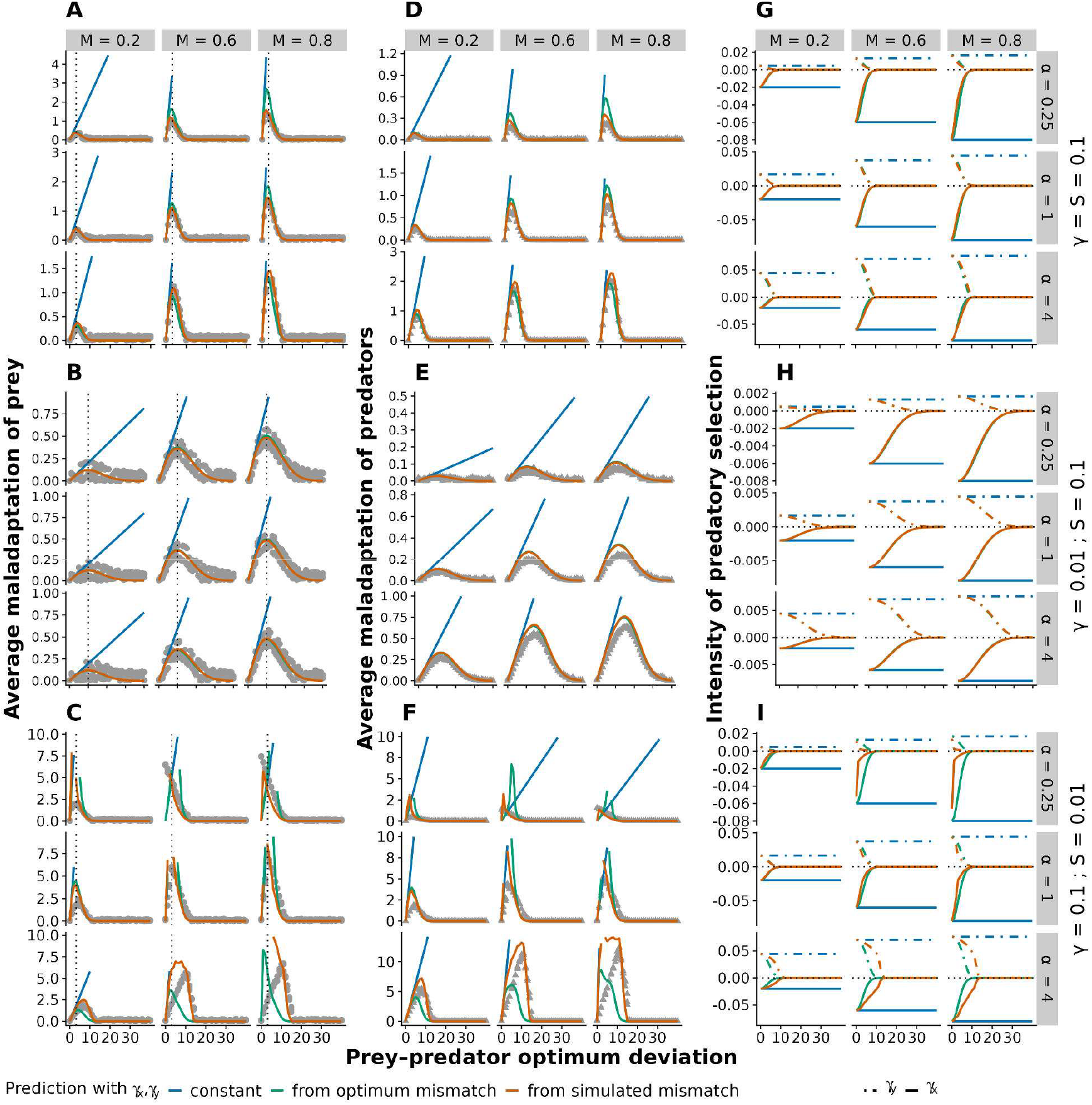
Influence of parameters of stabilizing and predatory selection in a constant environment with constant population sizes. The phenotypic deviation from optimum in prey (A-C) and predators (D-F), and the intensity of predatory selection *γ*_*x*_ in prey and *γ*_*y*_ in predators (G-I), are plotted against the difference between the optimum phenotypes in prey and predators, for a range of combinations of the strength of stabilizing selection *S*_*x*_ *= S*_*y*_ *= S* and selectivity of the interaction *γ*. All representations are as in Figures 2B (for panels A-C), S1 (panels G-I), and S2 (panels D-F), and all other parameters are as in these figures.

**Figure S4:**
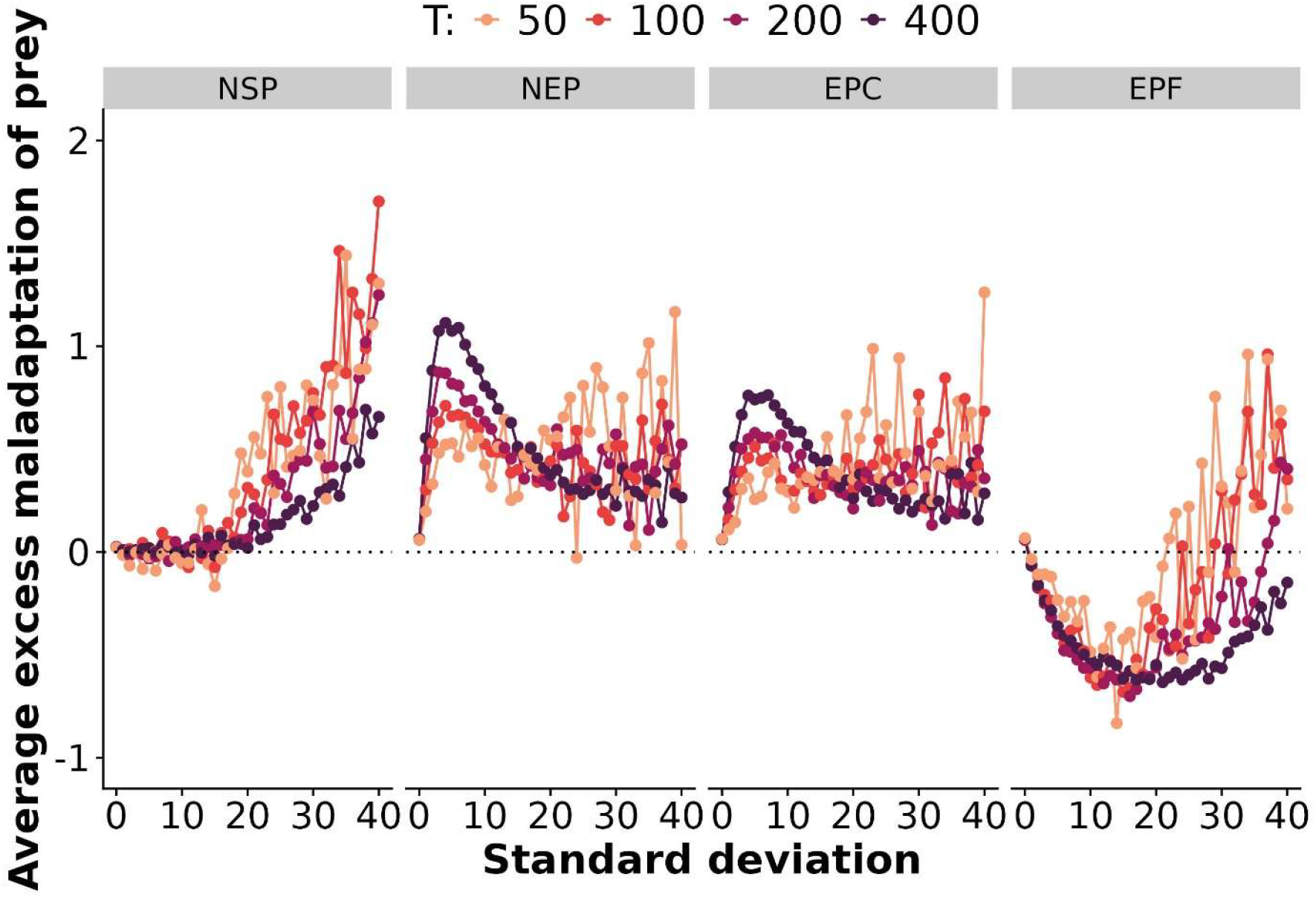
Adaptive tracking in a stochastically fluctuating environment with constant population sizes. The standard deviation of prey phenotypic deviation from optimum, shown as difference with the no-predator (NP) scenario (referred to as excess maladaptation), is represented against the standard deviation of fluctuations in the optimum, for different characteristic times of fluctuations *T* (colors), as explained below eq. (16) in the Methods. Results are averaged over 500*T* generations after a burn-in period of 5*T* generations, conducted under a Wright–Fisher model. Parameters are *γ = 0*.*01; N = 10000; P = 10000; S*_*x*_ *= 0*.*01; S*_*y*_ *= 0*.*01* and *T* as indicated in the legends. Four scenarios regarding predators are represented: a non-selective predator (NSP), a non-evolving predator (NEP), an evolving predator with a constant optimum at 0 (EPC), and an evolving predator with the same fluctuating optimum as the prey one (EPF).

**Figure S5:**
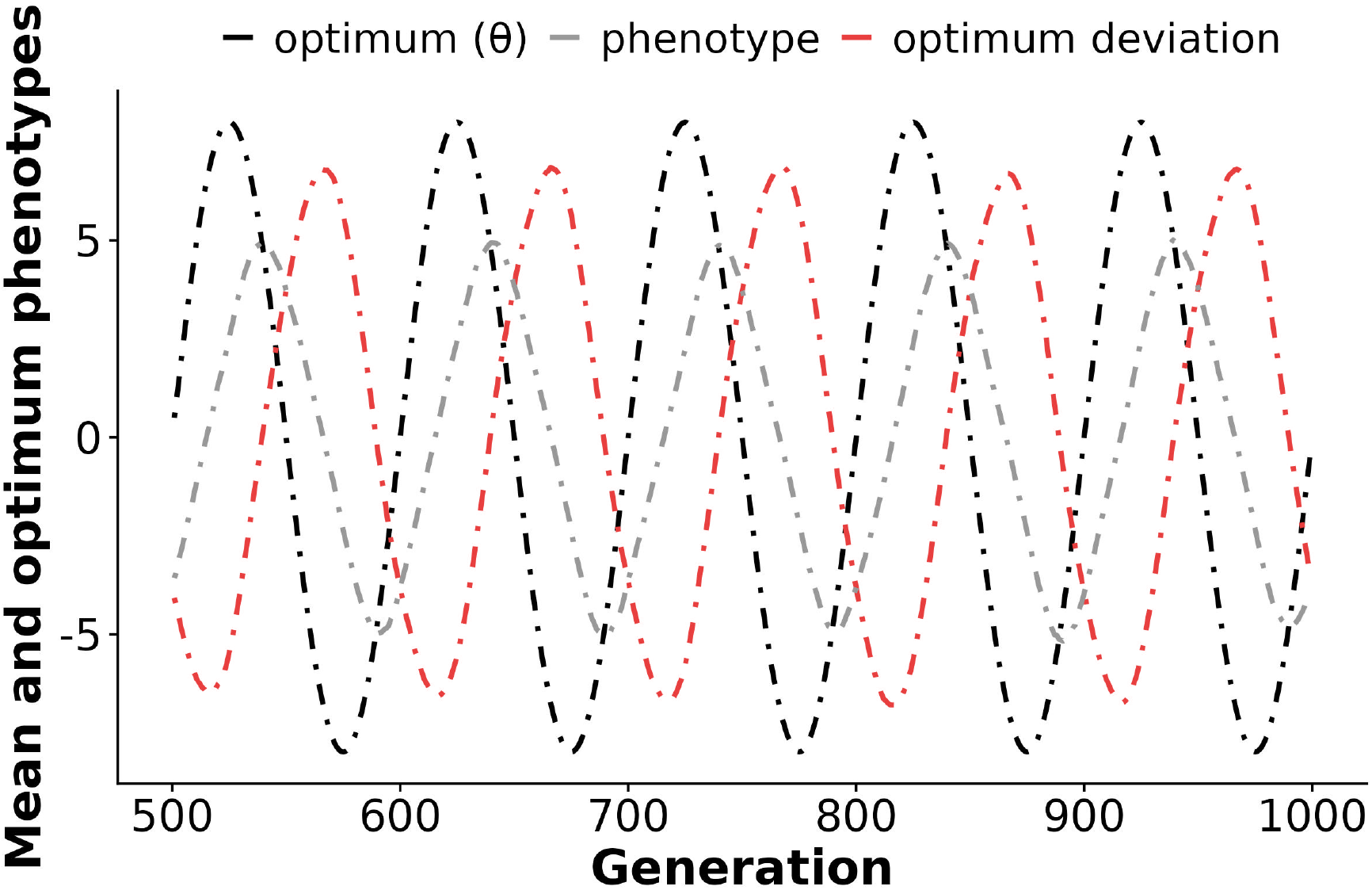
Evolution of the predator in a periodically fluctuating environment with constant population size. The mean phenotype of predators is shown in gray, for 500 generations of simulations of a Wright–Fisher population adapting to a periodically fluctuating optimum (shown in black), in a scenario where a selective predator tracks the same fluctuating optimum as the prey (EPF). The deviation of the predator’s mean phenotype from its optimum is also plotted in orange. Parameters are *γ = 0*.*01; T = 100; A = 8; N = 10000; P = 10000; α = 1; M = 0*.*6; S*_*x*_ *= 0*.*01; S*_*y*_ *= 0*.*01*.

**Figure S6:**
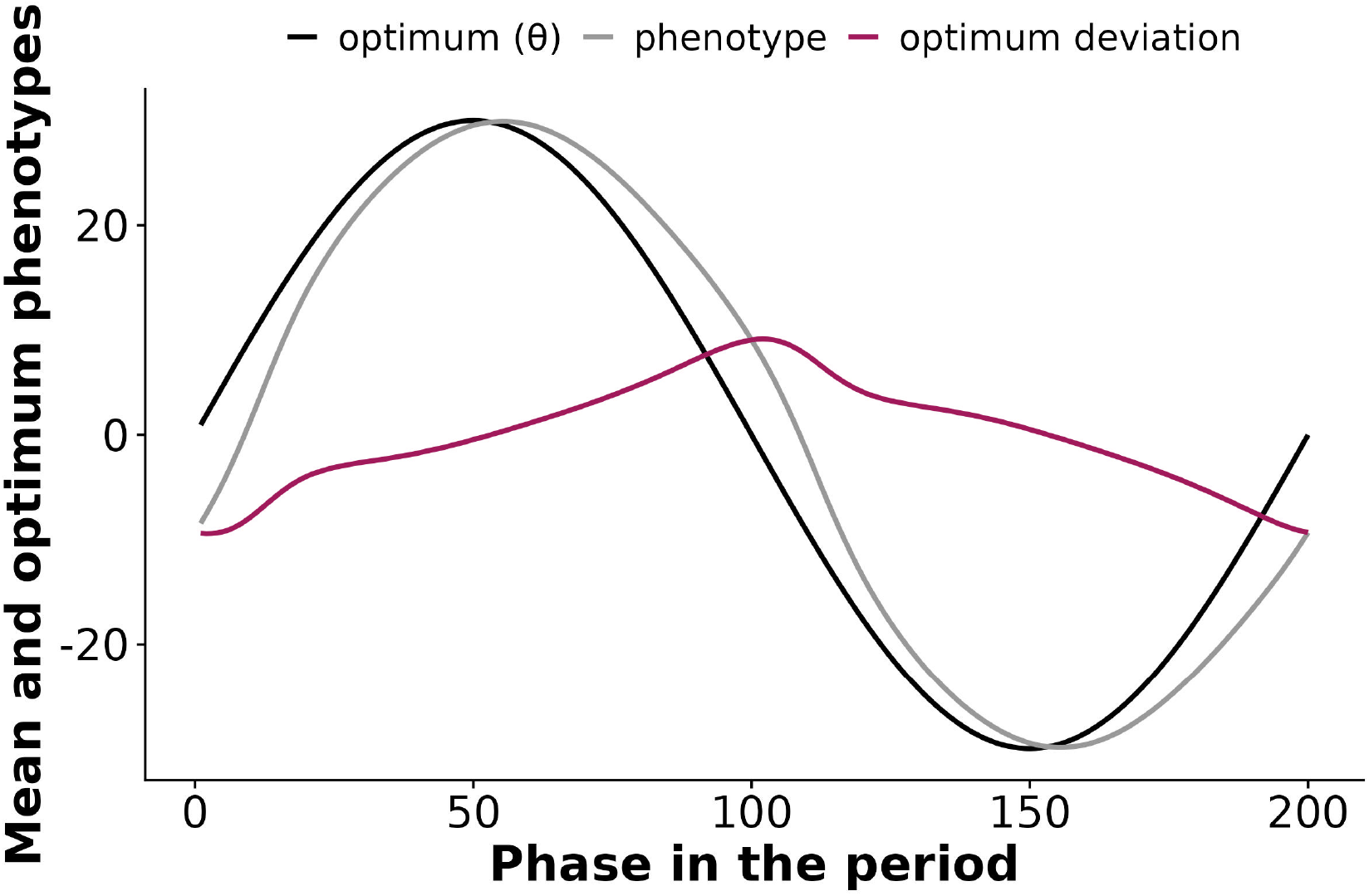
Illustration of an extreme case where prey deviation from the optimum deviates from a sine wave. The mean phenotype of prey (gray), the optimum phenotype (black), and the deviation of the mean phenotype from the optimum (purple), are shown for Wright–Fisher simulations of prey and predators, in a scenario where the optimum for prey fluctuates periodically, while the (evolving) predator has a constant optimum at 0 (EPC). Values were average over 100 phased periods, after a burn-in 5 periods. Parameters are: *γ = 0*.*01; T = 200; A = 30; N = 10000; P = 10000; α = 1; M = 0*.*6; S*_*x*_ *= 0*.*01; S*_*y*_ *= 0*.*01*.

**Figure S7:**
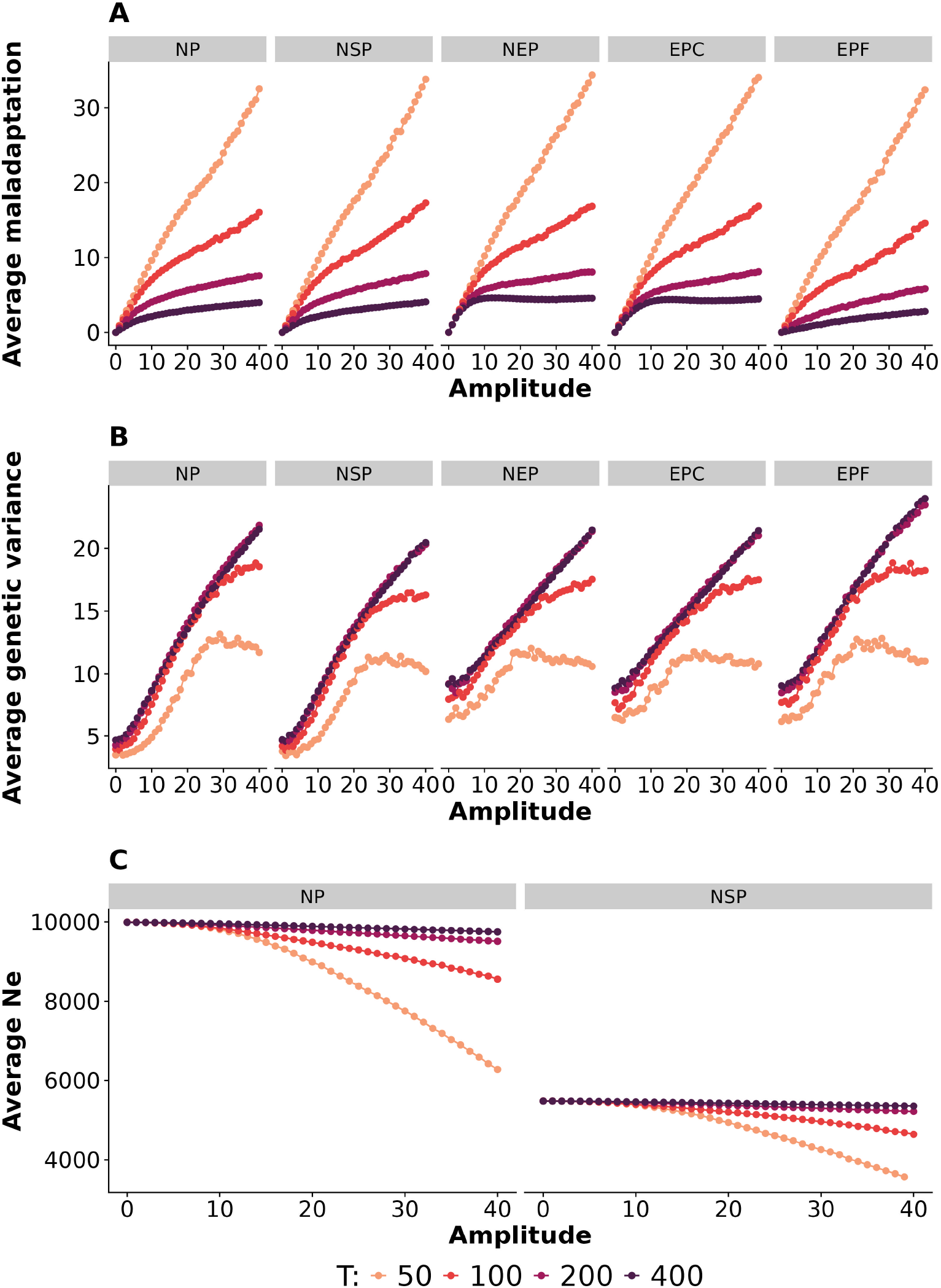
Adaptive tracking in a periodically fluctuating environment with constant population sizes. A) Amplitude of prey phenotypic deviation from optimum and B) Mean genetic variance of prey as a function of the amplitude of the optimum fluctuation. The results are based on the same simulations as in Figure 3 (BC) in the main text, except that values have not be normalized by taking the difference with scenario NP. C) Harmonic mean effective population size in simulations of 100 periods after a burn-in period of 5 periods, conducted under a Wright–Fisher model for prey and predator populations, across a range of amplitude of fluctuation (from 0 to 40). Parameters are as in panel A. Two scenarios have been investigated: no predators (NP), and non-selective predators (NSP).

**Figure S8:**
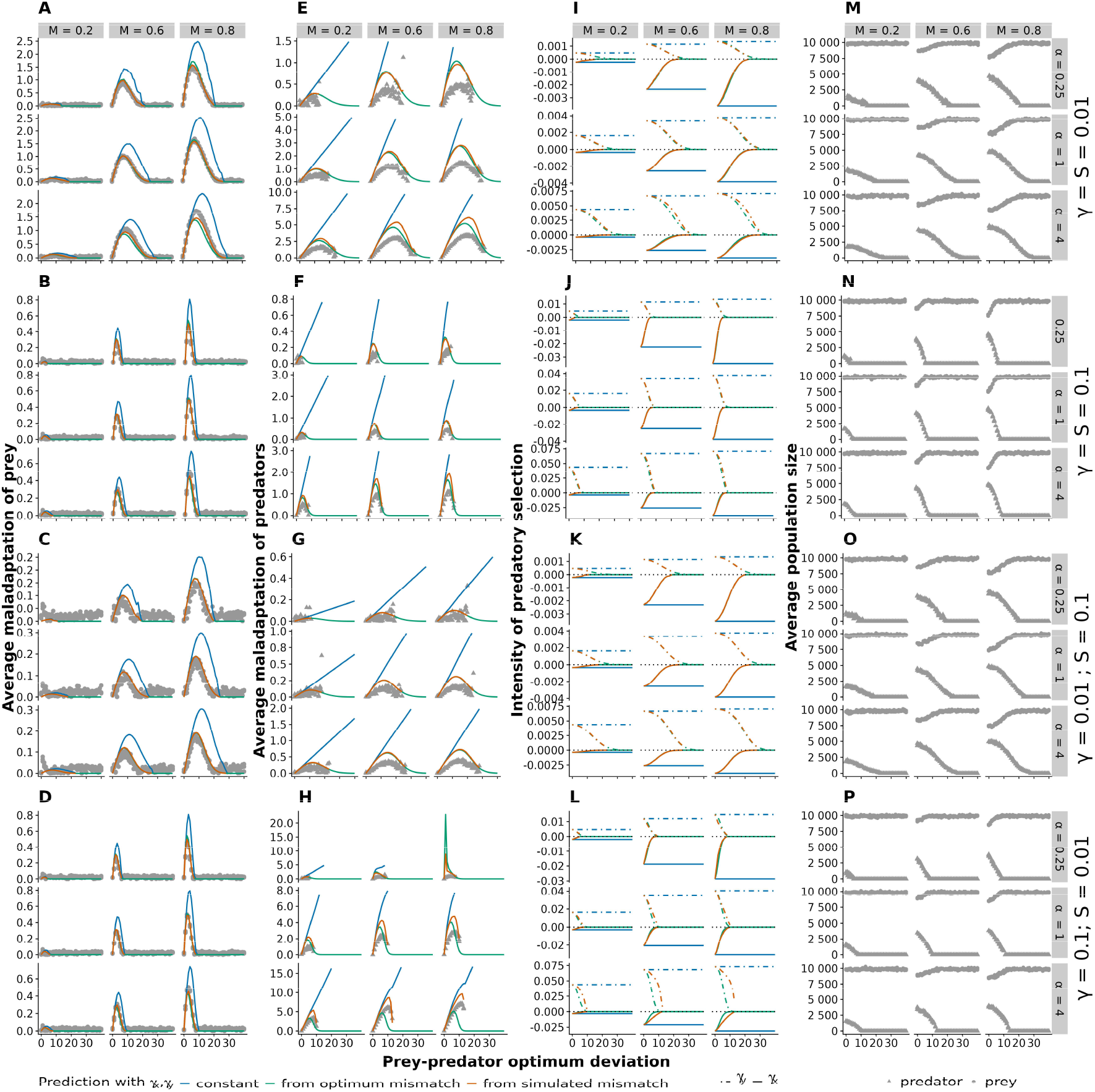
Influence of parameters of stabilizing and predatory selection in a constant environment with eco-evolutionary dynamics. The first three columns are similar to Figure S3 but conducted with eco-evolutionary dynamics rather than with Wright-Fisher populations. The last column (M-P) represents the population sizes of prey (gray dots) and predators (gray triangles). Missing dots in panels A-H correspond to simulations in which the predator population got extinct.

**Figure S9:**
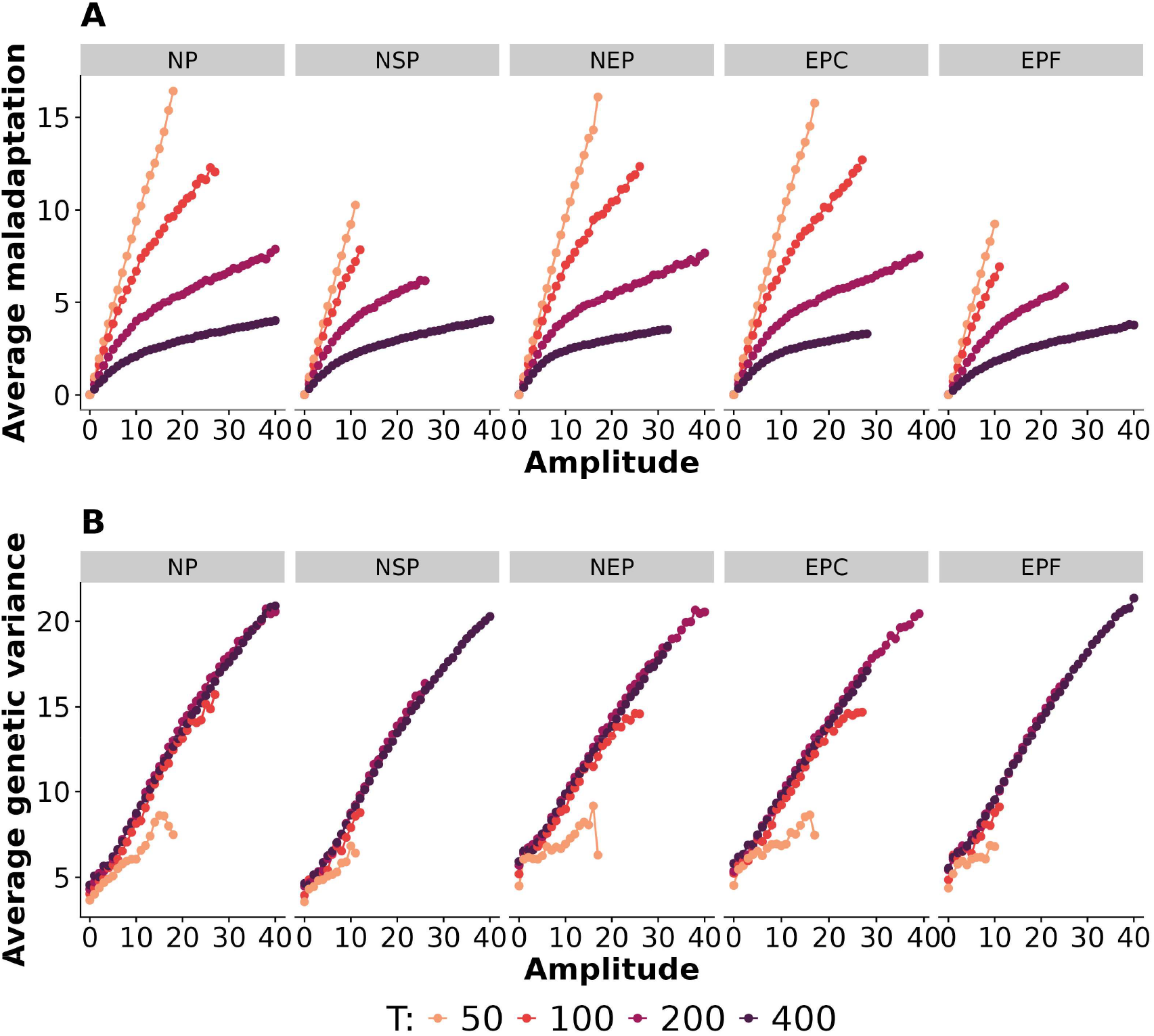
Adaptive tracking in a periodically fluctuating environment with eco-evolutionary dynamics. A) Amplitude of prey phenotypic deviation from optimum and B) mean genetic variance of prey as a function of the amplitude of the optimum fluctuation. Results are shown from simulations of 100 periods after a burn-in period of 5 periods, conducted for prey and predator populations with eco-evolutionary dynamics, across a range of amplitudes of fluctuation (from 0 to 40). Simulations were performed with *γ = 0*.*01; S*_*x*_ *= 0*.*01; S*_*y*_ *= 0*.*01; α = 1; M = 0*.*6* and *T* as indicated. Five scenarios regarding predators are represented: a no-predator scenario (NP), a non-selective predator (NSP), a non-evolving predator (NEP), an evolving predator with a constant optimum at 0 (EPC), and an evolving predator with the same fluctuating optimum as the prey one (EPF). Missing dots correspond to simulations in which prey or predator population got extinct.

